# Structure-guided development of a potent human B^0^AT1 inhibitor effective in a mouse model of phenylketonuria

**DOI:** 10.1101/2025.10.27.684712

**Authors:** Takuya Imazu, Tomoya Akashi, Masahiro Hiraizumi, Yosuke Inui, Wataru Sasaki, Tsuyoshi Takahashi, Hidenori Todoroki, Taichi Kumanomidou, Kazunori Yamada, Norie Fujikawa, Hiromi Hisano, Hidetsugu Asada, Tsukasa Kusakizako, Tomohiro Nishizawa, So Iwata, Osamu Nureki, Ikuko Miyaguchi

## Abstract

B^0^AT1 is a neutral amino acid transporter responsible for the (re)absorption in the intestine and kidney. Here, we developed small-molecule inhibitors of B^0^AT1 as a therapeutic strategy to a transient pocket, located ∼17 Å away from the substrate-binding site and unique to the outward-open conformation, and stabilize an outward-occluded conformation that prevents the conformational transitions required for transport. Guided by structural insights, we optimized an initial inhibitor (Cinromide) to improve potency and cross-species activity, yielding compound 3, which inhibits both human and mouse B^0^AT1 with submicromolar IC_50 values. In a PKU mouse model (*Pah^enu2^*), oral administration of compound 3 increased urinary Phe excretion and significantly reduced plasma Phe. Our findings identify a druggable allosteric site in B^0^AT1, demonstrate its utility for achieving potent and selective inhibition in vivo, and establish allosteric blockade as a therapeutic approach for PKU and other SLC6-family transporters.

## Introduction

Phenylketonuria (PKU) is a recessive genetic disorder caused by deficiency of phenylalanine hydroxylase (PAH), leading to phenylalanine (Phe) accumulation and severe neurodevelopmental impairment. Elevated Phe disrupts brain amino acid homeostasis, resulting in hypomyelination, gliosis, and neurotransmitter deficits^1,2^.

Although drugs like sapropterin and pegvaliase are approved for PKU, they are ineffective in many cases, due to genetic heterogeneity. Thus, there remains a substantial unmet need for broadly effective therapies to lower blood Phe levels across diverse patient populations^3^.

We targeted B^0^AT1 to reduce the plasma Phe levels in patients with PKU.

B^0^AT1 (SLC6A19) is a key transporter that mediates absorption of neutral amino acids, including Phe, in the intestine and reabsorption in the kidney. Inhibition of B^0^AT1 function in the kidney is expected to block Phe reabsorption into the blood, promote its urinary excretion, and lower its plasma levels^4–6^. B^0^AT1 is a member of the SLC6 transporter family, which includes serotonin transporter (SERT; SLC6A4), IMINO transporter (SIT1; SLC6A20), and dopamine transporter (DAT1; SLC6A3)^7–9^. The activity of B^0^AT1 depends on the co-expression of Tmem27 (Collectrin) in the kidneys and angiotensin-converting enzyme 2 (ACE2) in the intestines^10^. Similar dependencies are observed for other neutral amino acid transporters, such as B^0^AT3 and SIT1 ^10,11^. The ACE2–B^0^AT1 complex is also known as a receptor for coronaviruses^12^. Co-expression of human ACE2 or Collectrin is known to increase transporter function by approximately tenfold^13,14^.

B^0^AT1 has a leucine transporter-like (LeuT) fold structure common to SLC5, 6, and 7, with ten α-helices in TM1-10. The function and structure of this transporter family have been extensively studied in the LeuT and serotonin transporter (SERT) of SLC6^7,15^. The structure of LeuT consists of a bundle domain of four α-helices (TM1, 2, 6, and 7) and a hash domain of another four α-helices (TM3, 4, 8, and 9), with gate helices TM5 and 10. These transporter classes utilize sodium ions (Na^+^) as a co-substrate for active transport. Extracellular Na^+^ binding induces a conformational change in the hash domain, shifting the transporter to an outward-facing conformation. This allows for substrate binding, followed by a transition to an inward-facing conformation, simultaneously translocating both the substrate and Na^+^ into the cell. The stoichiometry of Na^+^ binding varies, ranging from one to three ions, and some transporters also require chloride ions (Cl^-^) for function. This cyclical process, characterized by alternating outward and inward conformations, is known as the alternating access mechanism^16^. Although the amino acids at the Na1 and Na2 sites are commonly found in SLC6s, as previously reported^16^, B0AT1 transports substrates in a 1:1 ratio with Na⁺, independent of Cl⁻^17,18^. Therefore, the number of Na^+^ sites and their roles remain unclear.

Several B^0^AT1–ACE2 structures^19,20,8,21^ have been reported, revealing the substrate-binding S1 site adjacent to the conserved unwound regions in TM1 and TM6, which exhibits structural flexibility that favors the recognition of multiple substrates^22^. Although an allosteric binding mode for Cinromide analogs was recently proposed, its functional role had not been fully elucidated. Previous cryo-EM studies have revealed that the allosteric site located between TM6 and TM8 and stabilize the outward-open conformation to prevent the transition to the inward conformation^21,23,24^. Importantly, this site is distinct from both the canonical substrate-binding site (S1) and the S2 site of SERT, which is often referred to as an allosteric site and accommodates inhibitors like escitalopram and duloxetine^25–27^. Cinromide analogs bind to a unique, previously uncharacterized site specific to B^0^AT1, topologically separate from any known ligand-binding pockets in other SLC6 transporters.

Contrary to the limited size and sequence conservation in the S1 site, allosteric sites offer a more promising avenue for developing specific inhibitors. However, *in vivo* efficacy for B^0^AT1 allosteric inhibitors has yet to be reported. Successful drug development necessitates demonstrating efficacy in animal models and humans, bridging the gap between in vitro and in vivo studies. This requires iterative compound optimization to achieve target protein efficacy across species.

In this study, we developed inhibitors that specifically bind to an allosteric site in B^0^AT1 and demonstrated that these inhibitors reduce plasma Phe concentrations *in vivo* in PKU model mice. Cryo-EM structures of multiple conformations revealed that this site is exclusive to the outward-open conformation and that binding of the compounds induces an outward-occluded conformation without substrate binding. These insights pave the way for the development of therapeutics targeting this allosteric site for the treatment of PKU.

## RESULTS

### Sample preparation and structural characterization of B0AT1 in apo and Cinromide-bound forms

Human wild-type B^0^AT1 (encoded by *SLC6A19*) was unstable when expressed alone. To address this, we stabilized the protein through complex formation with ACE2 and Collectrin or by introducing stabilizing mutations and antibody Fab fragments. We selected Cinromide (compound 1; Fig 1a), a known potent B^0^AT1 inhibitor^23,28,29^, as an initial compound for structural analysis.

**Figure 1.**
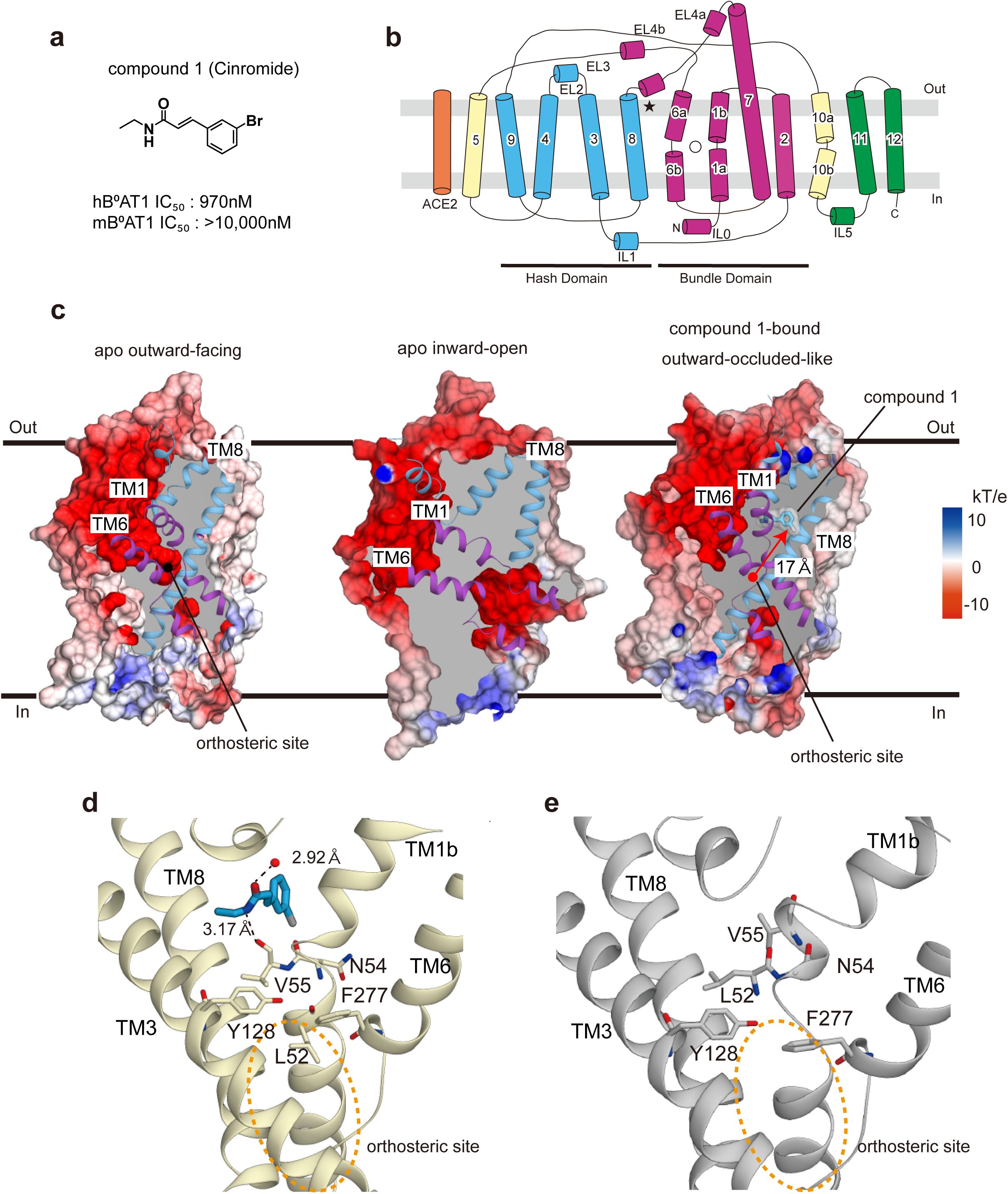
Cryo-EM structures of the hB^0^AT1 in the three states. a) Structure of compound 1 (Cinromide), a B⁰AT1 inhibitor, with IC₅₀ values for human and mouse B⁰AT1. b) Topological diagram of the hB⁰AT1-ACE2 heterodimer, showing the location of the orthosteric (〇) and compound binding (★) sites. c) Cross-sectional electrostatic surface potentials of apo outward-facing B⁰AT1, apo inward-open B⁰AT1 and the hB⁰AT1–compound 1 complex (outward-occluded-like). d) The compound binding site and its surrounding structure in the hB⁰AT1-compound 1 complex. e) The same site of apo outward-facing hB⁰AT1. The gate formed by Y128 and F277 is conserved in both structures.

Cryo-EM analysis of B⁰AT1 in complex with ACE2 or Fab yielded three distinct conformations: an outward-facing apo structure (ACE2 complex), an inward-open apo structure (Fab complex), and an outward-occluded-like closed structure bound to compound 1 (ACE2 complex). A topological overview of the B⁰AT1–ACE2 complex is shown in Fig. 1b. To provide a visual comparison of the three conformations, we generated cross-sectional views of the structures, shown with electrostatic surface potentials in Fig. 1c. The outward-facing apo structure was consistent with previous reports^19^. Notably, we identified a sodium ion (Na^+^) in the Na2 site, coordinated by five neighboring atoms (Extended Data Fig. 1a). In contrast, the inward-open apo structure lacked a density for Na⁺ at the Na2 site (Extended Data Fig. 1b). No electron density was detected at the putative Na1 site in either apo structure (Extended Data Fig. 1c, d), likely reflecting the absence of substrate^17^.

The compound1-bound structure adopts an outward-occluded-like conformation, with a closed intracellular gate formed by Tyr128 and Phe277, in the absence of substrate (Fig. 1d). In this conformation, Na⁺ was not observed at either the Na1 or Na2 sites (Extended Data Fig. 2a, b).

Compound 1-bound to an extracellular allosteric site ∼17 Å from the substrate-binding site (Fig. 1c, d). Its binding induced a shift of TM1b toward the orthosteric site, enabling a hydrogen bond between the amide NH of compound 1 and the carbonyl oxygen of Val55. The amide carbonyl of compound 1 also binds to a water molecule (Fig. 1d, Extended Data Fig. 2c). For comparison, the apo outward-facing structure is shown in Fig. 1e. In the compound 1–bound structure, Leu52 flipped toward the substrate-binding site, effectively occluding the cavity. In the previously reported structure of the inhibitor JX98, a hydrogen bond was formed with Trp56, and a similar Leu52 flipping was observed. However, unlike compound 1, it lacked the water-mediated interaction (Extended Data Fig. 3).

For compound 1, consistent with its binding mode, we observed potent inhibition of human B^0^AT1 (IC₅₀ = 970 nM), whereas inhibition of mouse B^0^AT1 was much weaker (>10,000 nM) (Fig. 1a).

### Optimized compounds and their improved in vitro activities

Because of the low inhibitory activity of compound 1 against mouse B^0^AT1(mB^0^AT1), *in vivo* efficacy testing in mice is not feasible. Therefore, we optimized compound 1 to improve its inhibitory potency against mB^0^AT1 (Fig. 2a-d). Our strategy focused on modifying interactions with two residues near the compound 1 binding site: Ile136 and Val317 in human B^0^AT1, which correspond to Val136 and Ala317 in mB^0^AT1 (Extended Data Fig. 4a, b). We hypothesized that these species-specific amino acid differences underlie the poor inhibition of mouse B^0^AT1 by compound 1.

**Figure 2.**
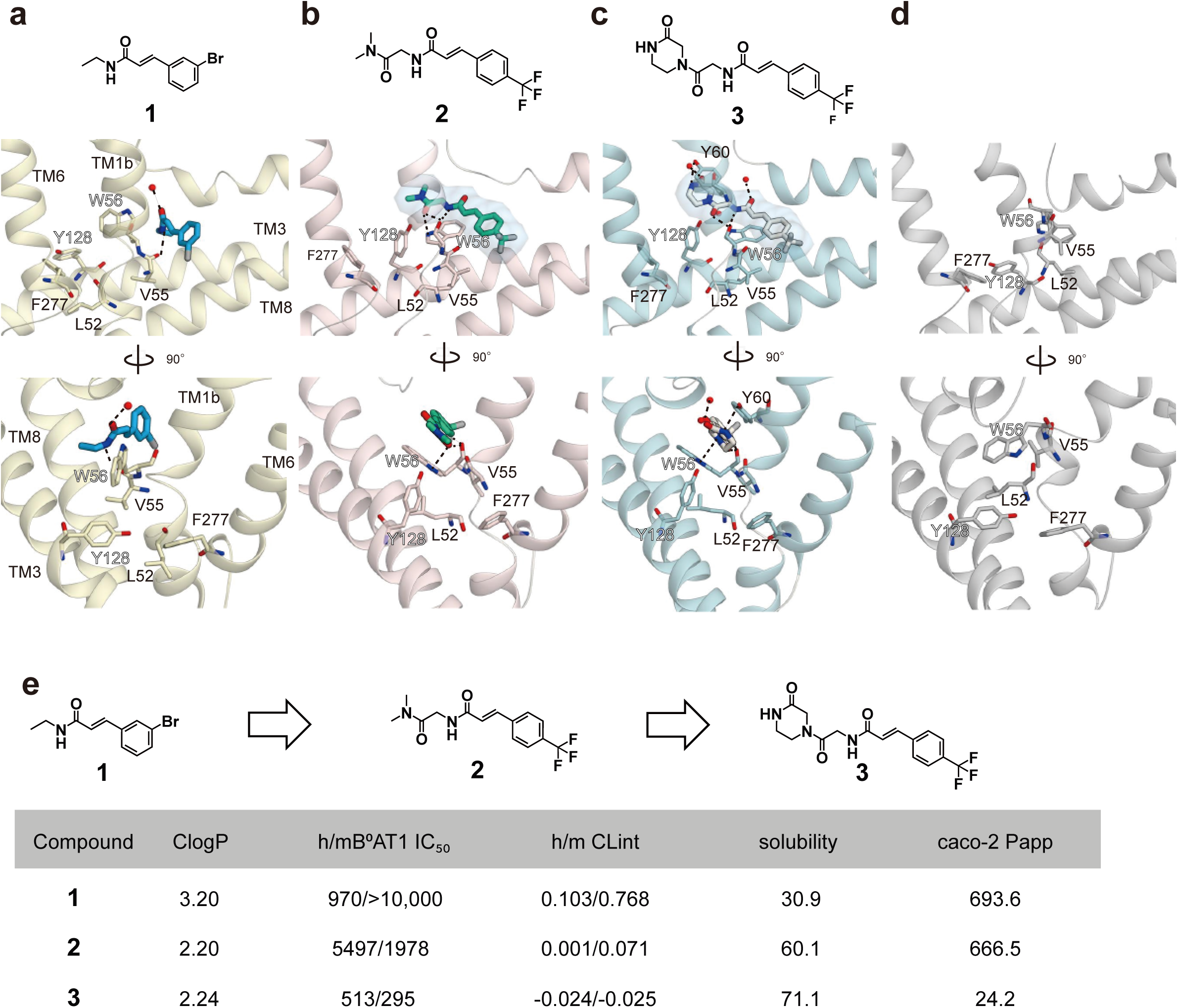
Comparison of binding sites and properties of compounds 1–3. a–c) Cryo-EM structures of B⁰AT1 bound to compound 1 (a), compound 2 (b), and compound 3 (c). Compounds 2 and 3, unlike compound 1, form interactions with Trp56 and occupy the cavity between TM1b and TM6 near the orthosteric site. d) The same binding site in the outward-facing apo structure. e) Comparison of inhibitory activities against human and mouse B⁰AT1(IC_50_, nM), along with metabolic stability (Clint, mL/min/g protein), membrane permeability (Papp, × 10^-7^ cm/sec), lipophilicity (ClogP), and solubility (JP2, µg/mL).

Ile136 is located near the ethylamine terminus of compound 1, and Val317 is near the cinnamic acid terminus. We reasoned that alkyl interactions might be insufficient for high-affinity binding to mB^0^AT1. To enhance binding and improve metabolic stability, we introduced chemical modifications, including a para-CF₃ group on the cinnamic acid ring and replacement of the ethylamine with a dimethylaminoglycine group. The resulting compound 2 showed improved metabolic stability and reduced lipophilicity, but at the expense of potency against human B⁰AT1 (IC50 for hB^0^AT1/ mB^0^AT1 are 5497 nM/1978 nM, respectively; Fig. 2e).

Cryo-EM structures revealed that compound 2 binds to the same extracellular allosteric site as compound 1, with Na⁺ absent from the canonical sodium-binding sites (Fig. 2a, b, Extended Data Fig. 5a-c). Unexpectedly, compound 2 induces local rearrangements in the surrounding amino acids.

Specifically, compound 2 forms hydrogen bonds with the main chain carbonyl oxygen of Trp56, which shifts toward TM3 and forms π interactions with the compound’s amide group (Fig. 2b). As a result, the Leu52 side chain reverted to its apo-like conformation, occluding the substrate-binding cavity (See Extended Data Fig. 9 for a structural comparison of the occluded and non-occluded conformations). Because compound 2 successfully reduced the species difference and gained inhibitory activity against mouse B^0^AT1, we further improved its potency by engaging the flexible regions of TM1 and TM6 and occupying the orthosteric-adjacent cavity. To this end, we replaced the dimethylamino group of compound 2 with a piperazinone moiety. The resulting compound 3 exhibited improved potency against both hB^0^AT1 and mB^0^AT1, with minimal species difference (IC₅₀: 513 nM for hB⁰AT1, 295 nM for mB⁰AT1; Fig. 2e). The cryo-EM structure showed that the piperazinone group extends closer to the substrate-binding pocket while maintaining a binding mode similar to that of compound 2 (Fig. 2c, Extended Data Fig. 5d-f).

Cell-based binding assays using ACE2–B⁰AT1 co-expressing cells revealed that the binding affinities (Kd) of compounds 1–3 were 472, 140, and 47.0 nM, respectively (Extended Data Fig. 6). Thermal stability assays showed increases of 1.1, 2.7, and 6.4 °C, respectively, relative to the control (44.5 °C) (Extended Data Fig. 7). Notably binding affinity (Kd) does not exactly much with their inhibitory potency (IC50); compound 2 exhibited lower inhibitory potency than compound 1. These findings indicate that compound 1 may induce more substantial conformational changes upon binding, contributing to its inhibitory effect.

Finally, to further investigate the inhibition mechanism of compound 3, we performed kinetic assays using radiolabeled d5-Phe uptake in cells expressing Collectrin–B⁰AT1. Compound 3 increased the Km for d5-Phe without affecting V_max_, indicating competitive inhibition (Fig. 3a, b, Supplementary Data Table 1). In addition, binding assays in the presence and absence of Na^+^ revealed a higher B_max_ value under Na⁺-rich conditions indicating that compound 3 preferentially binds to the Na⁺-bound conformation of B⁰AT1 (Fig. 3c, Supplementary Data Table 2). Structural comparison of the Na2 site in the apo and inhibitor-bound conformations further showed that compound 3 binds near the Na2 site without directly coordinating the ion (Fig. 3d, e).

**Figure 3.**
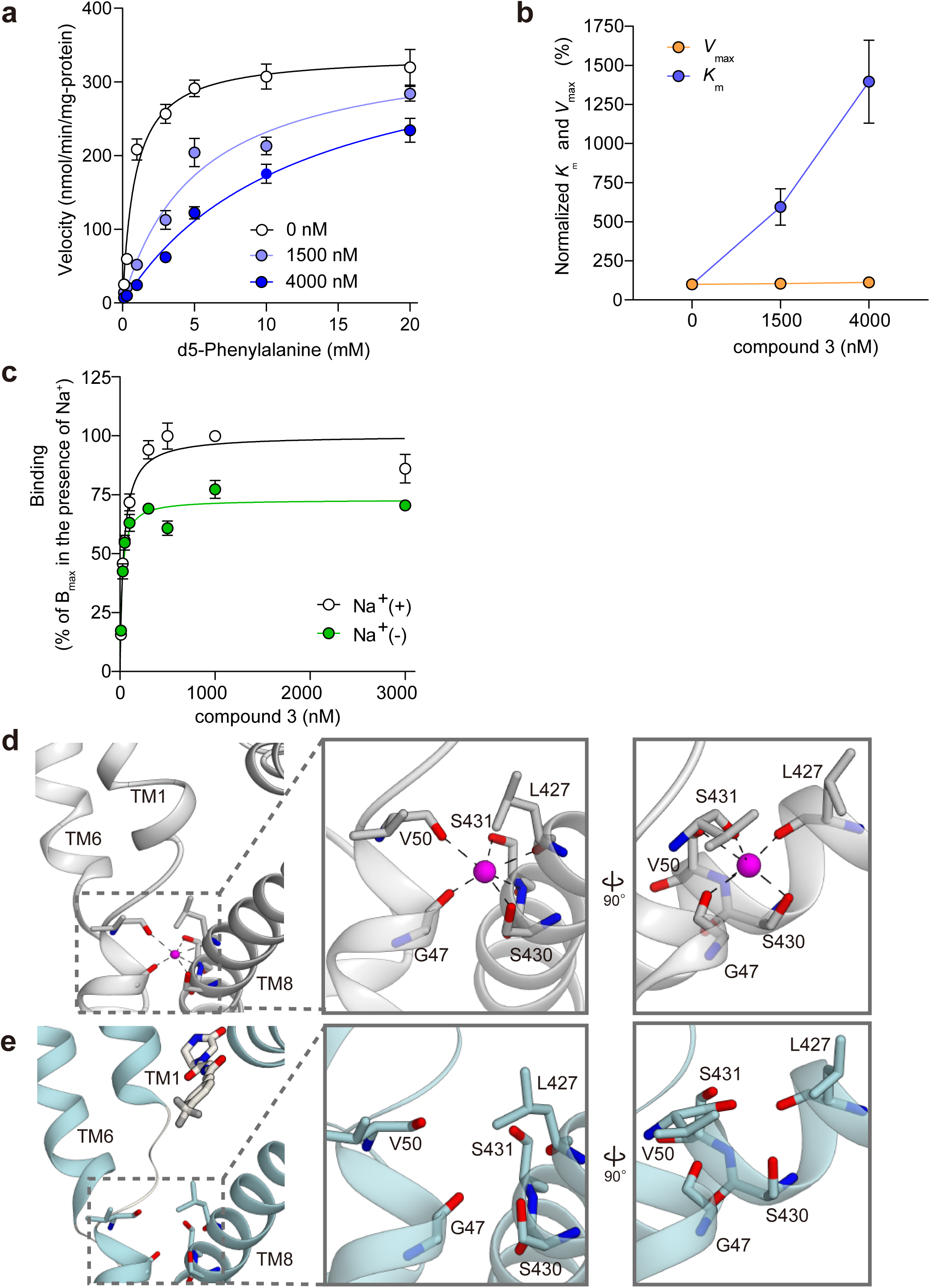
Competitive binding of B^0^AT1 inhibitors. a) Competitive inhibition of compound 3 in d5-phenylalanine uptake by hB⁰AT1. Each point represents mean ± SEM (n = 3, biological replicates). b) Kinetics parameters in d5-phenylalanine uptake by hB^0^AT1 in the presence of compound 3. Each point represents mean ± SEM (n = 3, biological replicates). c) Concentration-dependent binding of compound 3 in the presence and absence of Na^+^. Specific binding was normalized to the Bmax in the presence of Na^+^. Each point represents mean ± SEM (n = 3, technical replicates). d) Na2 Na^+^ ion binding site in the apo structure. e) Complex structure of compound 3 with B^0^AT1 around the Na2 Na^+^ ion binding site.

### In vivo pharmacological effects in Pah^enu2^ mice

Because of its favorable metabolic stability and solubility, we evaluated the *in vivo* effect of compound 3 on phenylalanine (Phe) levels in *Pah^enu2^* mice (Fig. 4a)^30^. These mice carry a homozygous F263S mutation in the Pah gene, resulting in a markedly reduction in PAH enzymatic activity and elevated plasma Phe levels, and detectable urinary excretion. They serve as a well-established model of PKU. Following oral administration of compound 3 (30 mg/kg), the compound was detectable in plasma at 4 and 8 hours post-dose (Supplementary Data Table 3). Although compound 3 shows low membrane permeability (24.2 × 10^-7^ cm/s; Fig. 2e), its detectable plasma levels are likely attributable to its relatively low molecular weight and high thermodynamic solubility.

**Figure 4.**
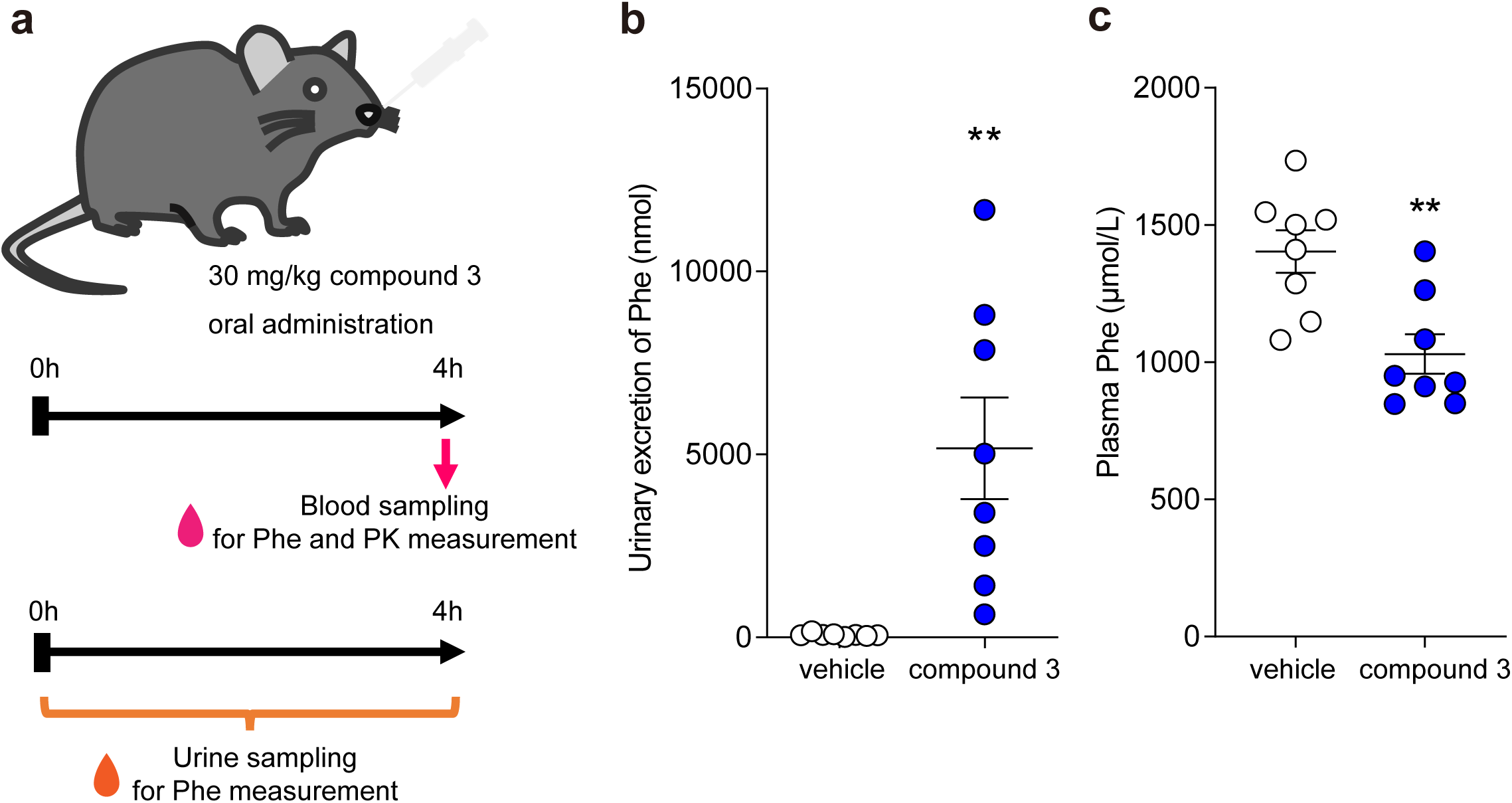
In vivo Pharmacological Effects of compound 3 in *Pah^enu2^* Mice. a) Schematic of the in vivo study design. *Pah^enu2^* mice were orally administered compound 3 (30 mg/kg), and blood and urine samples were collected after 4 hours post-dose. b) Urinary excretion of phenylalanine (Phe) measured 4 hours after compound 3 administration. Mice were placed in metabolic cages immediately after dosing, and urine was collected over 4 hours. c) Plasma Phe concentrations measured 4 hours after oral administration of compound 3. Phe levels were quantitated by LC–MS/MS. Data are shown as mean ± s.e.m. (n = 8 mice per group). Each dot represents an individual mouse. **p < 0.01 compared to vehicle group (Student’s t-test)

In 12-week-old male *Pah^enu2^* mice, compound 3 treatment significantly increased urinary Phe excretion and concomitantly reduced plasma Phe concentrations 4 hours after dosing (Fig. 4b, c). These findings are consistent with the proposed mechanism of B^0^AT1 inhibition, whereby reduced intestinal Phe absorption promotes urinary excretion and lowers systemic Phe levels.

#### Structural and functional analyses of the inhibition mechanism

We next examined the inhibition mechanism of the developed B^0^AT1 inhibitors by comparing the present cryo-EM structures with related transporter conformations.

B^0^AT1 functions through an alternating-access mechanism, in which Na⁺ and substrate bind simultaneously to the outward-facing state, triggering transitions through occluded intermediates to the inward-open conformation (Extended Data Fig. 8).

In the compound-bound structures, Compound 1 disrupted the substrate pocket by flipping Leu52, whereas Compounds 2 and 3 stabilized an occluded conformation lacking substrate (Extended Data Fig. 9). Functional assays showed that Compound 3 bound more strongly in the presence of Na⁺ and competed with Phe (Fig. 3a–c; Supplementary Data Tables 1, 2).

Comparison of the outward-open, outward-facing, and inward-open structures indicated that the inhibitor pocket exists only in the outward-open state (Extended Data Fig. 8b). MD simulations of LeuT indicate that Na^+^ binding at the Na2 site induces the transition from inward-open to outward-facing and enhances extracellular substrate binding^31,32^.

Consistent with this, Na⁺ binding at the Na2 site was observed in the outward-facing apo structure but not in the inward-open conformation (Extended Data Fig. 1a, b). Although no Na⁺ density appeared in the inhibitor-bound structures, binding assays confirmed that Compound 3 associated with B^0^AT1 even without Na⁺.

Together, these observations indicate that Na⁺ is not essential for inhibitor binding but facilitates formation of the outward-open conformation, thereby promoting inhibitor access to the allosteric pocket.

## DISCUSSION

Based on the structural and functional analyses above, we propose a mechanism in which B⁰AT1 inhibitors exert their effect by binding to a transient pocket that appears in the outward-substrate-binding site, thereby preventing substrate amino acids from accessing the transporter and effectively halting its activity (Fig. 5).

**Figure 5.**
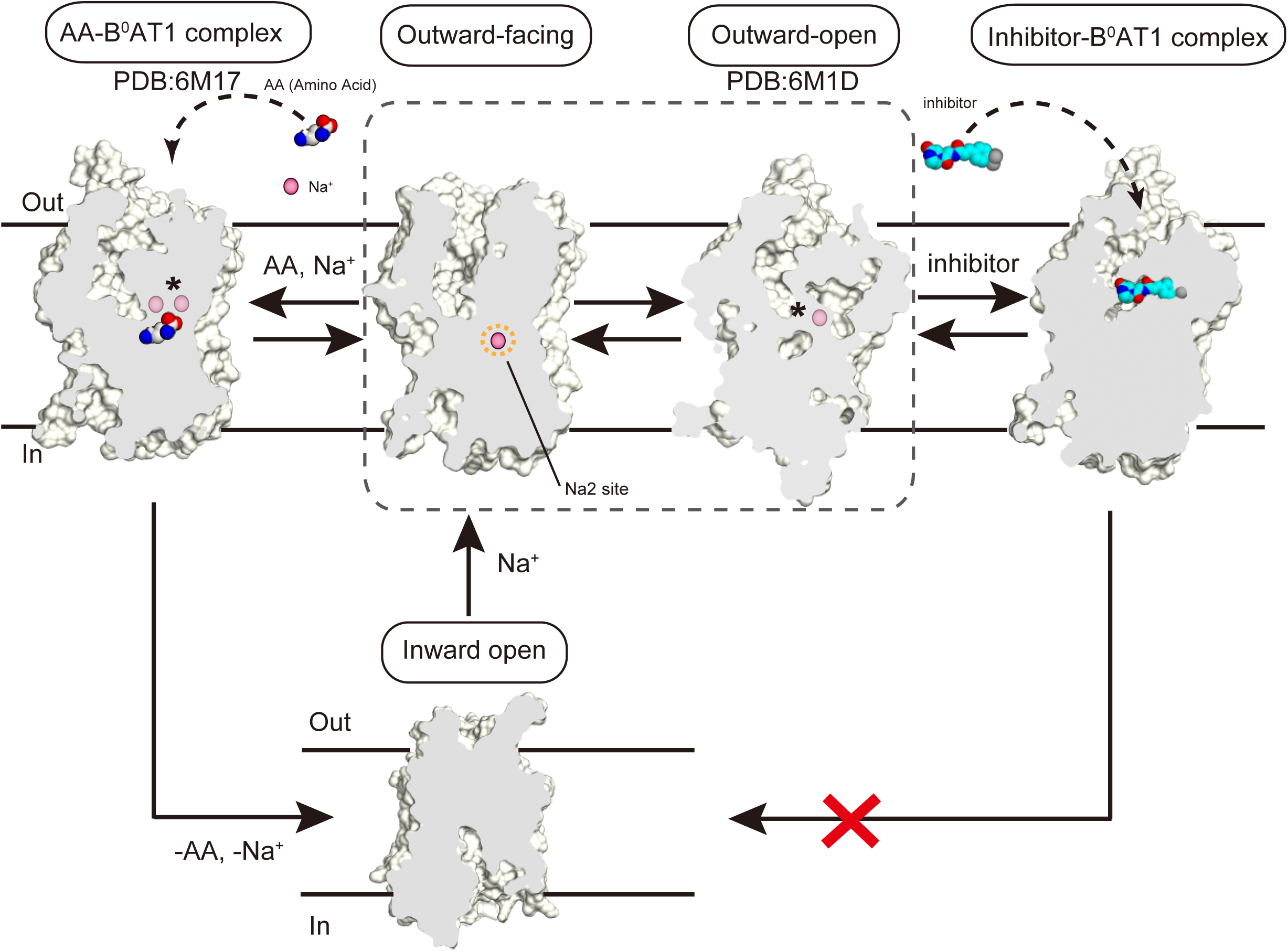
The inhibition mechanism of B⁰AT1 by Na^+^-coupled structural transitions. B⁰AT1 inhibitors bind to the outward-open conformation of B⁰AT1. When the B⁰AT1 inhibitor binds, the transport cycle stops. The asterisk indicates that it is not clear whether a Na^+^ is bound, how many Na^+^ are bound, or to which Na site they are bound. The figure shows the tentative number of Na^+^ assumed from the previous and following transport cycles. The outward-open conformation structure was generated using PDB:6M1D. For clarity, the surface representation was sectioned along two planes to highlight the continuous channel from the extracellular entrance to the central cavity. The AA–B⁰AT1 complex structure was generated using PDB:6M17.

Notably, this binding event stabilizes the outward-facing conformation and prevents the transition to the inward-facing conformation. As a result, the compound-bound transporter is unable to proceed through the transport cycle. This mechanism supports a model in which the inhibitors function by locking the transporter in an inactive, outward-facing conformation^21^.

In addition, the inhibitors exhibited competitive inhibition despite binding to an allosteric site distinct from the canonical substrate-binding pocket (Fig. 3a, b). This indicates that allosteric inhibitors can functionally mimic competitive inhibition by inducing conformational occlusion of the substrate-binding site.

Although compound 2 bound more tightly to B^0^AT1 than compound 1, it exhibited weaker inhibition in functional assays (Fig. 2e; Extended Data Fig. 6d, e; Extended Data Fig. 7). This apparent discrepancy indicates that binding affinity or thermal stability alone does not ensure effective inhibition. Instead, the ability of an inhibitor to induce conformational changes that restrict substrate access appears to be the key determinant of potency. Structural analysis supports this notion: compound 1 induces more pronounced rearrangements around the extracellular gate, effectively occluding the substrate-binding cavity, whereas compound 2 causes only localized adjustments. Thus, the degree of conformational locking, rather than affinity per se, underlies the observed difference in inhibitory activity.

To better understand the functional significance of the allosteric inhibition observed in this study, we next examined the structural characteristics of the compound-binding pocket. This pocket is located at the outer gate of B^0^AT1 and is formed by transmembrane helices TM1b, TM3, TM7, and TM8— core elements of the LeuT-fold scaffold that are involved in conformational transitions (Extended Data Fig. 9).

In transporter-targeted drug design, orthostatic binding sites typically serve as the primary targets. However, becauseB⁰AT1 transports a broad range of small amino acids, designing substrate analogues with high specificity is challenging. In contrast, the allosteric site identified here provides a promising alternative: it is structurally stable, suitable sized for small-molecule binding, and functionally essential. While the overall architecture of this site is conserved, sequence divergence at key positions may allow for selective recognition by small molecules. (Extended Data Fig. 10a).

Notably, this allosteric site is also present, albeit more restricted, in the bacterial transporter LeuT (Extended Data Fig. 10b; PDB: 3TT1). Moreover, a recent study reported an allosteric inhibitor binding to the equivalent site in the dopamine transporter DAT, reinforcing the idea that this pocket may serve as a generalizable target for drug development across structurally related transporters.

Together, these features highlight the allosteric pocket as a structurally and pharmacologically tractable site for the development of selective B^0^AT1 inhibitors. Compound 1 showed inhibitory activity against human B^0^AT1 but not against mouse B^0^AT1, whereas compounds 2 and 3 showed no species-specific differences. Compounds 2 and 3 likely share similar binding modes between the two species, as supported by comparable inhibitory potencies and binding affinities (Extended Data Fig. 6). In contrast, compound 1 bound only to human B^0^AT1, despite the high sequence identity (86.8%) between human and mouse orthologs. Although Val317 and Ile136—residues differing between species near the binding site (Extended Data Fig. 5)—were initially considered as potential contributors to the observed species difference, they do not fully account for the loss of compound 1 activity in mouse B^0^AT1. The structure of mouse B^0^AT1 has not been reported, but given the overall sequence similarity, the binding pocket is expected to be largely conserved.

One possible explanation lies in the dynamic nature of transporters: substrate binding and transport involve large-scale conformational changes, and species-specific differences outside the binding site may influence these structural transitions and affect compound efficacy.

These findings underscore the importance of considering not only static binding pockets but also conformational flexibility and remote residue contributions when designing inhibitors targeting structurally dynamic transporters like B^0^AT1.

Starting from compounds with in vitro inhibitory activity against B^0^AT1, we developed potent inhibitors that reduce plasma phenylalanine levels in a PKU mouse model. These inhibitors act via a newly identified allosteric site, offering high specificity and the potential for reduced side effects— key features for the lifelong management of PKU. Notably, we found that extracellular Na⁺ stabilizes the outward conformation of B^0^AT1 and enhances the binding of allosteric inhibitors, which is advantageous for in vivo use given the high Na⁺ concentration in plasma.

Overall, this study demonstrates that targeting non-orthosteric, conformationally dynamic allosteric sites can yield selective and effective inhibitors, providing a promising strategy for drug discovery beyond conventional substrate-binding approaches.

## ONLINE METHODS

### Reagents and chemicals

The synthesis of compounds 2 and 3 is described in the Supplementary Information. Compound 1 was purchased from Angene International, Ltd. (Nanjing, China). Phenylalanine was purchased from Sigma-Aldrich (St. Louis, MO). D5-phenylalanine was purchased from C/D/N Isotopes (Quebec, Canada).

### cDNA constructs

Wild-type-B^0^AT1 (UniProt accession: Q695T7) and ACE2 (UniProt accession: Q9BYF1) constructs were prepared as previously described^13^. B^0^AT1 complementary DNA and human ACE2 cDNA were synthesized and codon-optimized for expression in human cell lines. Both cDNAs were respectively subcloned into the pcDNA3.4 vector. An N-terminal FLAG tag was fused to B^0^AT1, and a Strep-tag was fused after the N-terminal signal peptide of ACE2.

### Expression and purification of wild-type B^0^AT1–ACE2

Mammalian FreeStyle 293-F cells (ThermoFisher Scientific, Waltham, USA) were grown and maintained in FreeStyle 293 Expression Medium (ThermoFisher Scientific, Warrington, USA) at 37 °C and 8% CO_2_ under humidified conditions. The cells were transiently transfected at a density of 1.8 × 10^6^ cells/mL with the plasmids and PEI Max (Polysciences). For transfection, approximately 270 μg of the B^0^AT1 plasmid and 270 μg of the ACE2 plasmid were premixed with 1890 μg of PEI Max in 60 mL medium for 15–30 min before transfection. Then, 60 mL of this mixture was added to 540 mL of the cell culture (600 mL total) and incubated at 37 °C in the presence of 8% CO_2_ for 72 h before collection. The cells were collected by centrifugation (1000 ×*g*, 10 min, 4 °C) and stored at - 80 °C before use. To prepare the B^0^AT1–ACE2 complex, the harvested cells were solubilized for 2 h at 4 °C in solubilization buffer (25 mM Tris-HCl [pH 8.0], 150 mM NaCl, 1% GDN [Anatrace], and protease inhibitor cocktail). After ultracentrifugation (20,000 ×*g*, 45 min, 4 °C) and filtering through a syringe filter (0.45 μm pore size), the supernatant was incubated with M2 Anti-FLAG Affinity Resin (Sigma-Aldrich) equilibrated with buffer A (25 mM Tris-HCl [pH 8.0], 150 mM NaCl, 0.02% GDN) and incubated for 2 h at 4 °C. The resin was washed with 5 column volumes (CVs) of buffer A, and the protein was eluted with buffer A supplemented with 0.1 mg/mL DYKDDDDK Peptide (Fujifilm Wako, Tokyo, Japan). The eluate was mixed with Strep-Tactin Sepharose (IBA), equilibrated with buffer A, and incubated for 1 h at 4 °C. After incubation, the resin was washed with 5CV buffer A, and the protein was eluted with buffer A plus 4 mM of desthiobiotin (Sigma-Aldrich). The sample was concentrated and purified via size-exclusion (Sigma-Aldrich) chromatography on Superose 6 Increase 10/300 GL (GE Healthcare) pre-equilibrated with the same buffer. The peak fractions were pooled and concentrated to 10–12 mg/mL.

### Design of stabilized B^0^AT1

As wild-type B^0^AT1 is prone to aggregation, stabilized B^0^AT1 was designed. Introduction of N-terminal and C-terminal truncates of B^0^AT1, substitution of surface-exposed cysteine with alanine, and point mutations were tested. Sequential alignment was performed with SLC6A family transporters (dopamine transporter [SLC6A3] and SERT [SLC6A4]), whose structures have been previously reported^33–35^, to introduce point mutations. Stabilized B^0^AT1 (hereinafter, sB^0^AT1) was successfully obtained by combining mutations (⊿2-7, Δ621-634, C500A, C610A, E81L, L114I, S290A, F471L, Y507W, and M542L), which contributed to the stabilization or improvement of expression, as assessed by fluorescence-detected size-exclusion chromatography (FSEC). For sB^0^AT1, an enhanced green fluorescent protein sequence and 8× polyhistidine tag at the C terminus of sB^0^AT1 were retained after the tobacco etch virus (TEV) cleavage site. The aforementioned stabilizing mutations were introduced into this construct via site-directed mutagenesis.

### Expression and purification of sB^0^AT1

Mammalian HEK293S GnTI-cells (ATCC) were grown and maintained in FreeStyle 293 Expression Medium supplemented with 2% fetal bovine serum at 37 °C and 8% CO_2_ under humidified conditions. Cells were transiently transfected at a density of 2.0 × 10^6^ cells/mL using plasmids and FectoPRO (Polyplus, Illkirch, France). Approximately 200 μg of sB^0^AT1 plasmid was premixed with 200 μL of FectoPRO reagent in 80 mL of medium for 10–20 min before transfection. For transfection, 80 mL of the mixture was added to 0.72 L of the cell culture and incubated at 37 °C with 8% CO₂ for 72 h before harvesting. The cells were collected by centrifugation (800 ×*g*, 10 min, 4 °C) and stored at −80 °C before use.

To prepare sB^0^AT1, the cells were solubilized for 2 h at 4 °C in buffer (50 mM Tris-HCl [pH 7.5], 250 mM NaCl, 20% glycerol, 1.5% [w/v] n-Dodecyl-β-D-maltoside (DDM; Sigma-Aldrich), 0.15% 3β-Hydroxy-5-cholestene 3-hemisuccinate; 5-Cholesten-3β-ol 3-hemisuccinate (CHS; Sigma-Aldrich), protease inhibitor cocktail (cOmplete, Roche, Basel, Switzerland). After ultracentrifugation (142,400 ×*g*, 30 min, 4 °C), the supernatant was incubated with Affi-Gel 10 (Bio-Rad) coupled with a GFP-binding nanobody^36^ and incubated for 3 h at 4 °C. The resin was washed five times with five column volumes of wash buffer (50 mM Tris-HCl [pH 7.5], 300 mM NaCl, 20% glycerol, 20 mM imidazole [pH 7.5], 0.1% DDM [Anatrace], and 0.005% CHS) and suspended in TEV protease and incubated overnight to cleave the eGFP-His tag. The flow-through was pooled, and the cleaved eGFP-His tag was removed by passing through TALON metal affinity resin pre-equilibrated with wash buffer.

N-linked glycans were removed by adding homemade His-tagged PNGaseF to the sample after TALON metal affinity resin treatment. After removing PNGaseF by passing through a Ni-NTA superflow, the samples were concentrated and purified by size-exclusion chromatography on a HiLoad Superdex75 16/600 column (GE Healthcare) equilibrated with SEC buffer (20 mM Tris-HCl [pH 7.5], 150 mM NaCl, 0.05% DDM [GLYCON Biochemicals], and 0.005% CHS). Finally, for the cryo-EM trial, the SEC buffer was replaced with a buffer containing 20 mM Tris-HCl (pH 7.5), 150 mM NaCl, and 0.06% GDN, using a Superdex200 Increase 10/300 GL column.

### Preparation of monoclonal antibody (Fab#54)

All animal experiments conformed to the Guide for the Care and Use of Laboratory Animals of Japan and were approved by the Kyoto University Animal Experimentation Committee. sB^0^AT1 was used to prepare the monoclonal antibody (Fab#54). Construct design, expression, and purification were performed as described above. Purified sB^0^AT1 was reconstructed in liposomes composed of 10:1 egg phosphatidylcholine (Sigma-Aldrich) to adjuvant lipid A (Sigma-Aldrich) to enhance the immune response. MRL/lpr mice were immunized with three injections of liposomal sB^0^AT1 at 2-week intervals. Spleen cells were collected from immunized mice and fused with myeloma cells. To select antibodies recognizing the sB^0^AT1 structure, multistep screening, including liposome-ELISA, denaturing ELISA, and fluorescence size-exclusion chromatography, was performed. For the cryo-

EM trial, a large amount of the selected monoclonal antibody against sB^0^AT1 (IgG #54) was prepared via a large-scale culture of hybridoma cells. Monoclonal antibodies were initially purified with protein G resin, eluted with Gly buffer, digested with papain immobilized on N-hydroxysuccinimide– Sepharose and incubated for 5 h at 37 °C. Protein A chromatography was used to separate the Fc gragment from the Fab. The collected Fab fragment was then purified by size-exclusion chromatography on a HiLoad Superdex200 16/600 column (GE Healthcare) equilibrated with SEC buffer (20 mM Tris-HCl [pH 7.5], 150 mM NaCl, 0.05% DDM [Anatrace], and 0.005% CHS). Finally, the SEC buffer was replaced with a buffer containing 20 mM Tris-HCl (pH 7.5), 150 mM NaCl, and 0.06% GDN by ultrafiltration (Amicon Ultra-15 30 K; Millipore). The sequence of Fab#54 was determined via standard 5’-RACE using total RNA isolated from hybridoma cells.

The sequence of the Fab#54 fragment light chain is provided below: AQAAELDIVMTQSPSSLVVSAGEKVTMTCKSSQSLFNSRTRKNYLAWYQQKPGQSPKLLIYWASTRESGVPDRFTGSGSGTDFTLTISSVQAEDLAVYYCKQSSYLLTFGAGTKLELKRADAAPTVSIFPPSSEQLTSGGASVVCFLNNFYPKDINVKWKIDGSERQNGVLNSWTDQDSKDSTYSMSSTLTLTKDEYERHNSYTCEATHKTSTSPIVKSFNRNEC The sequence of the Fab#54 fragment heavy chain is provided below: GGSSRSSLEVQLQQSGPELVKPGASVKISCKASGYTFTDYYINWVKQRPGQGLEWIGWIYPRSGGTKYIEKFKGKATLTVDTSSSTAYMQLNSLTSEDSAVYFCARFYYSYNEDYYALDYWSQGTSVTVSSAKTTAPSVYPLAPVCGDTSGSSVTLGCLVKGYFPEPVTLTWNSGSLSSGVHTFPAVLQSDLYTLSSSVTVTSSTWPSQSITCNVAHPASSTKVDKKIEPRGPTIKPCPPCKCPAPNLLGGPSVFIFPPKIKDVLMISLSPIVTCVVVDVSEDDPDVQISWFVNNVEVHTAQTQTHREDYNSTLRVVSALPIQHQDWMSGKEFKC

### Electron microscopy sample preparation

The purified wild-type B^0^AT1–ACE2 protein was incubated with compounds 1, 2, and 3 (0.5 mM) and incubated for 1 h on ice. The sB^0^AT1–Fab#54 antibody complex was assembled by incubating sB^0^AT1 with Fab#54 at a molar ratio of 1:1.8 overnight on ice. The mixture was subjected to size-exclusion chromatography (Superdex200 Increase 10/300 GL; GE Healthcare). Fractions containing the sB0AT1–Fab#54 complex were pooled and concentrated to 8–12 mg/mL using Amicon Ultra-15 centrifugal filters (30 kDa MWCO; Millipore). Immediately before grid preparation, 450 µM phenylalanine was added to the sB^0^AT1–Fab#54 sample.

Grids were glow-discharged in low-pressure air at 10 mA using a PIB-10 vacuum device (Mito, Japan). Samples—including compound-bound, apo, and B^0^AT1–antibody complexes—were applied to freshly glow-discharged Quantifoil R1.2/1.3 Cu/Rh 300-mesh holey carbon grids (SPT Labtech, Melbourn, UK) using a Vitrobot Mark IV (Thermo Fisher Scientific) at 4 °C, with a blotting time of 4–6 s under 99% humidity. Grids were then plunge-frozen in liquid ethane

### Electron microscopy data collection and processing

The sample grids were loaded onto a Titan Krios G3i transmission electron microscope (Thermo Fisher Scientific) operating at 300 kV, equipped with a Gatan Quantum-LS energy filter (GIF) and a Gatan K3 Summit direct electron detector in correlated double-sampling (CDS) mode. Images were acquired at a nominal magnification of 105,000×, yielding a calibrated pixel size of 0.83 Å/pixel, at the University of Tokyo, Japan.

#### For ACE2–B^0^AT1 samples

Movies were dose-fractionated into 48 frames at a dose rate of 13.0–13.7 e⁻/pixel/s in CDS mode, resulting in a total exposure of 49–51 e⁻/Å².

#### For antibody-bound sB^0^AT1

Movies were acquired in non-CDS mode with a dose rate of 7.57 e⁻/pixel/s over 48 frames, resulting in a total exposure of 52 e⁻/Å².

Data collection was automated using the image-shift method in SerialEM software, with a defocus range of −0.8 to −1.6 μm. The total number of images is listed in Table 1.

**Table 1.**
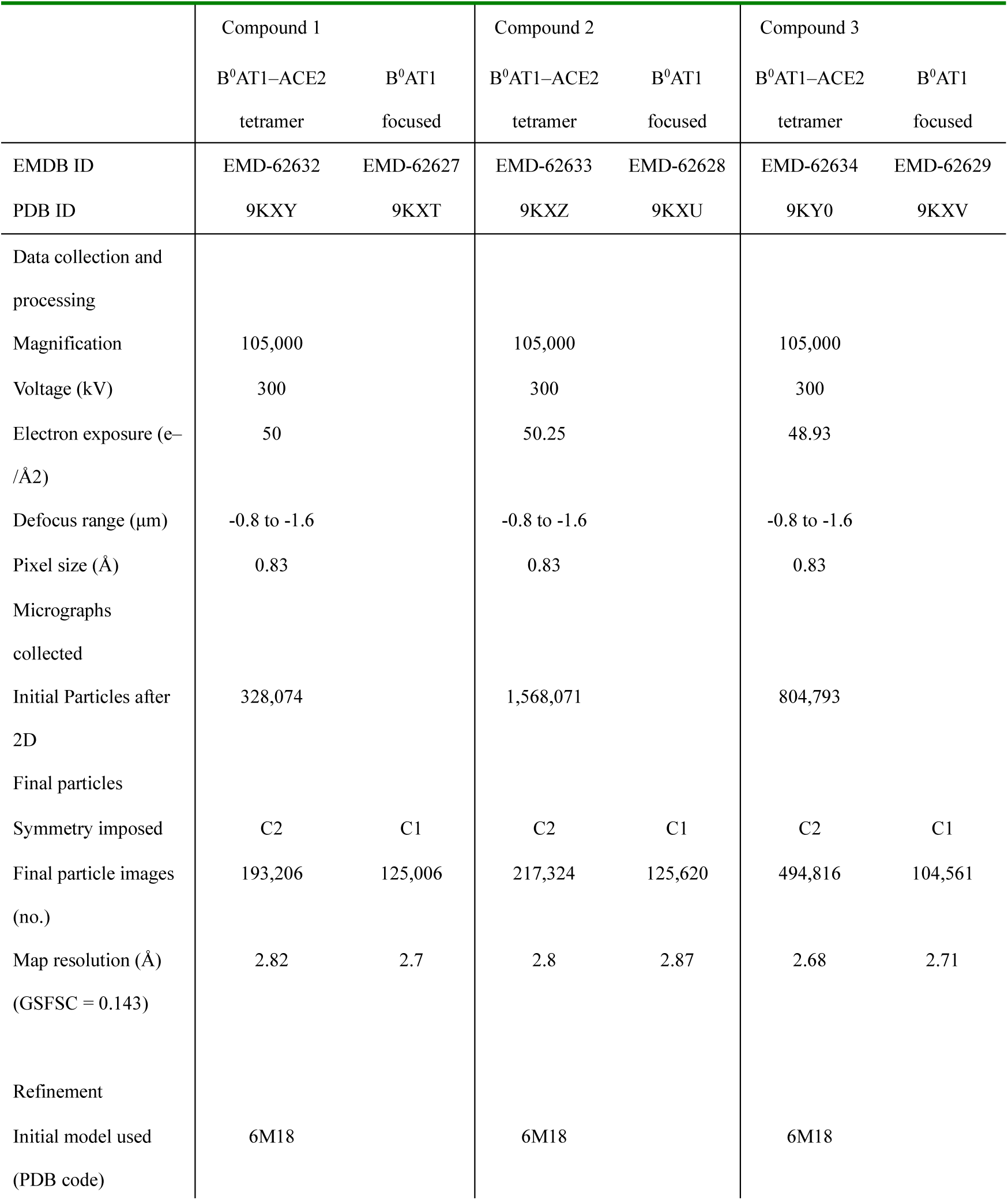
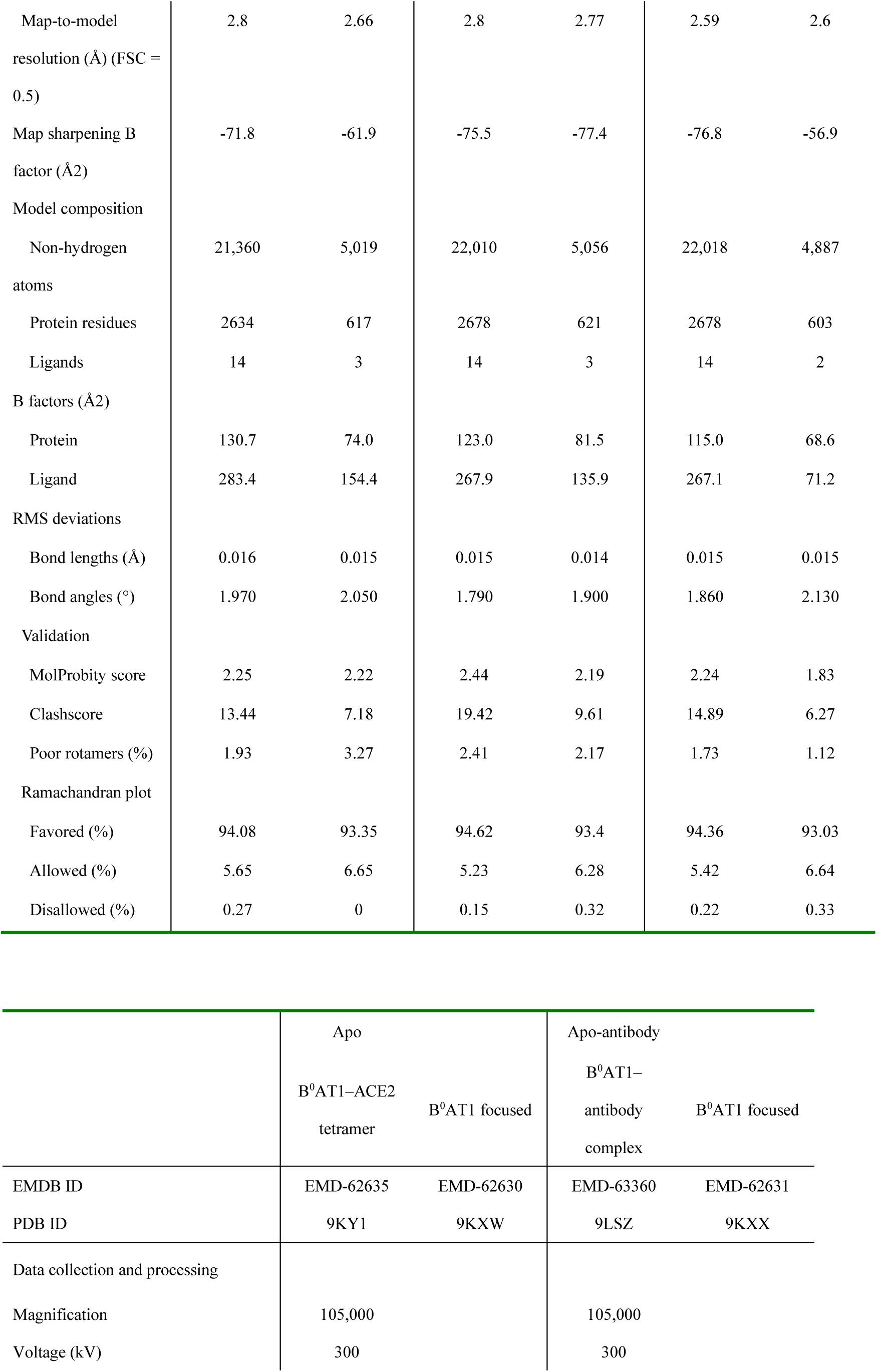
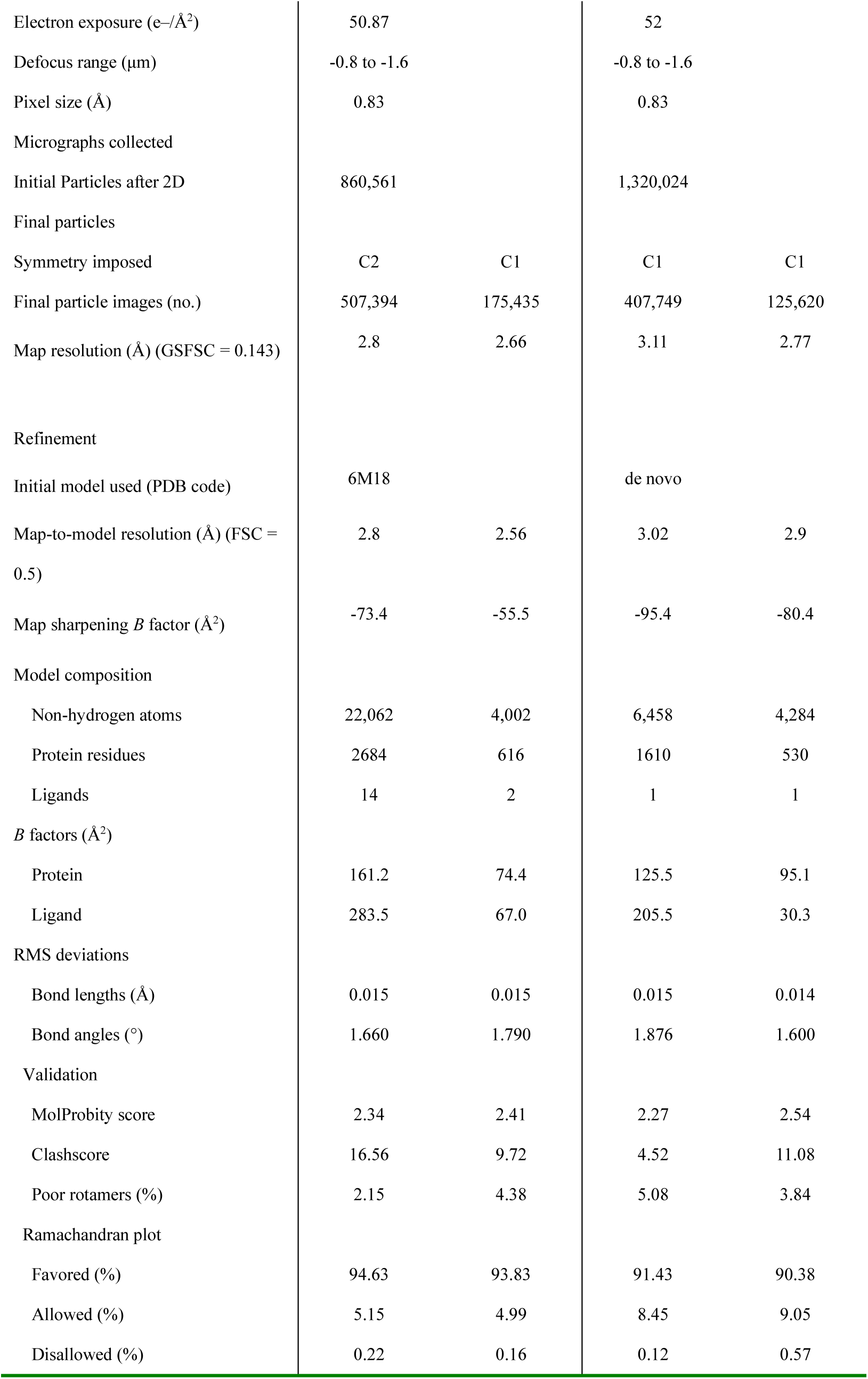
Cryoelectron microscopy (cryo-EM) data collection, refinement, and validation statistics.

For all datasets, dose-fractionated movies were subjected to beam-induced motion correction using RELION^37^, and contrast transfer function parameters were estimated using CTFFIND^38^.

For the compound 1–bound dataset, 1,310,641 particles were initially picked from 2,364 micrographs using the Topaz picking function in RELION. Particles were extracted by downsampling to a pixel size of 3.2 Å/px and subjected to several rounds of 2D and 3D classifications. The best class contained 193,206 particles, which were re-extracted with a pixel size of 1.107 Å/px and subjected to 3D refinement, yielding a 3D reconstruction of the B^0^AT1–ACE2 dimer at a global resolution of 2.82 Å. For improved resolution, we used the symmetry expansion and focused classification in RELION were employed for a single subunit of the B^0^AT1–ACE2 transmembrane region complex. The second 3D classification yielded four map classes. The best class contained 125,006 particles, which were subjected to further 3D refinement, yielding a map with a global resolution of 2.7 Å, according to the FSC = 0.143 criterion. Finally, the local resolution was estimated using RELION-4. This processing strategy is illustrated in Supplementary Fig. 1

The data of compound 2–3-bound or apo ACE2–B^0^AT1 (Supplementary Figs. 2–4) were collected and analyzed in the same manner. For the antibody-bound apo-sB0AT1(Supplementary Fig. 5), 2,009,409 particles were initially selected from 3,924 micrographs using the topaz-picking function in RELION. Particles were extracted by downsampling to a pixel size of 3.2 Å/px and subjected to several rounds of 2D and 3D classifications. The best classes contained 407,749 particles, which were re-extracted with a pixel size of 0.83 Å/px subjected to further 3D refinement, yielding a map of the B^0^AT1–Fab complex with a global resolution of 3.00 Å. Subsequent micelle subtraction and nonaligned 3D classification using a mask (without micelles) resulted in three map classes. The best class contained 125,620 particles and was further refined and postprocessed, yielding a map with a global resolution of 2.77 Å, according to the FSC = 0.143 criterion. Finally, the local resolution was estimated using RELION-4. This processing strategy is illustrated in Supplementary Fig. 5.

### Model building and validation

For B^0^AT1–ACE2 structures of apo and compound-bound forms, the Protein Data Bank (PDB) model PDBID: 6M18 was the starting model. The model of the apo-inward conformation of B^0^AT1-Fab was manually constructed *de novo* using COOT^33^, guided by the inward-facing structure of SERT (PDB ID: 6DZZ) and an Alphafold2^34^ -predicted model of the Fab fragment. The density map corresponding to the phenylalanine added to the Fab-B^0^AT1 sample was not visible and therefore not modeled.

After manual adjustment, models were structurally refined using the Servalcat^35^ pipeline in REFMAC5^39^ and manual real-space refinement in COOT. The 3D reconstruction and model refinement statistics are summarized in Table 1. All structural models in this study were built based on B^0^AT1-focused maps (Supplementary Fig. 6, 7). All molecular graphics were prepared using CueMol (http://www.cuemol.org), PyMOL (http://www.pymol.org/pymol), and UCSF Chimera^40^.

### Cell lines

The Jump-In TI platform cells (hereafter referred to as JTREx-CHO cells*) were used to generate a cell line expressing human SLC6A19 and human Collectrin via R4 integrase-mediated DNA integration at a defined target site in accordance with the Jump-InTM TITM Gateway® Targeted Integration System user guide (www.thermofisher.com)*. JTREx-CHO host cells were co-transfected with pJTI R4 Int and the tetracycline-inducible expression vector pJTR-hB^0^AT1, which carry the hSLC6A19-T2A-hCollectrin synthetic construct (synthesized by GenWiz), using Lipofectamine3000 (ThermoFisher Scientific). Successful integration of the expression vectors at the R4 attP site conferred neomycin resistance to JTREx-CHO cells by cointegrating the EF1a promoter upstream of the promoterless neomycin resistance gene. Retargeted cells were selected for G418 sulfate resistance, cloned, and used for measurements. Mouse SLC6A19-Collectrin co-expressing cells were similarly established using the pJTR-mB^0^AT1 vector carrying the mSLC6A19-T2A-mCollectrin synthetic construct (synthesized by GenWiz) instead of the pJTR-hB^0^AT1.

### d5-Phenylalanine uptake in Collectrin-B^0^AT1 co-expressing cells

JTREx-CHO hB^0^AT1 cells were maintained in Dulbecco’s modified Eagle medium/F-12 (ThermoFisher Scientific) supplemented with 10% fetal bovine serum (HyClone), 0.1 mg/mL hygromycin B, and 0.5 mg/mL M G418 at 37 °C in a humidified 5% CO₂ atmosphere. Cells were seeded at 2.0 × 10^5^ cells/well in 24-well plates with or without 2.5 µM doxycycline and cultured for 24 h. The medium was removed, the cells were washed twice and pre-incubated with extracellular fluid buffer (100 mM NaCl, 2 mM KCl, 1.8 mM CaCl_2_, 1 mM MgSO_4_, and 5 mM HEPES; pH 7.4) at 37 °C for 60 min with or without inhibitor. After preincubation, uptake was initiated by adding d5-phenylalanine solution to a final concentration of 200 µM. Uptake was completed by removing the uptake buffer and washing thrice with ice-cold buffer, followed by solubilization in 1 N NaOH. Cell lysates were deproteinized by adding acetonitrile containing d8-phenylalanine as an internal standard. The d5-phenylalanine concentration was quantified by liquid chromatography-tandem mass spectrometry (LC–MS/MS) using the internal standard method. Cellular protein content was determined using a bicinchoninic acid protein assay (ThermoFisher Scientific). d5-Phenylalanine uptake was expressed as the cell-to-medium ratio (μL/mg-protein; the ratio of cellular concentration [pmol/mg-protein] to medium concentration [pmol/µL]).

For inhibition analysis, the cell-to-medium ratio of cells incubated without doxycycline was used as the background. Specific d5-phenylalanine uptake was calculated by subtracting this background from the total cell-to-medium ratio and normalizing to the uptake achieved without the inhibitor. Kinetic parameters were calculated by nonlinear regression using GraphPad Prism 8.4.3 (San Diego, CA, USA; Supplementary Fig. 8).

### Thermostability measurement

Detergent-solubilized proteins were analyzed via fluorescence-detection size-exclusion chromatography-based thermostability assay (FSEC-TS) to monitor intrinsic tryptophan fluorescence using an ACQUITY UPLC BEH SEC 4.6 mm × 150 mm column (Waters). Purified B^0^AT1–ACE2 was mixed with the compound dissolved in DMSO (or DMSO only as control) to a final concentration of 0.1 µM B^0^AT1–ACE2, 500 µM compound (or no compound), and 0.5% DMSO and incubated at 4 °C for 60 min. The samples were then aliquoted into PCR tubes, 45 µL each, and incubated at 4, 10, 20, 25, 30, 35, 40, 42.5, 45, 47.5, 50, 52.5, 55, 60, 65, and 70 °C for 10 min in a PCR Thermal Cycler SP (Takara Bio). The samples were centrifuged again (20,000 ×*g*, 60 min, 4 °C) to clear the lysate, and 3 μL of the supernatant was placed in the SEC column, pre-equilibrated with buffer containing 50 mM Tris, pH 8.0, 150 mM NaCl, and 0.02% GDN (Anatrace).

### B^0^AT1 inhibitor binding assay

B^0^AT1 inhibitor binding was measured according to a previously reported method with minor modifications^41^. FreeStyle 293-F cells were co-transfected with hB^0^AT1 and hACE2 plasmids. The cells were collected and disrupted by sonication in a hypotonic buffer (50 mM Tris-HCl [pH 7.5], 10 mM KCl, and a protease inhibitor cocktail). Cell debris was removed by centrifugation (2,000 ×*g*, 5 min, 4 °C). The membrane fraction was collected by ultracentrifugation (112,000 ×*g*, 30 min, 4 °C) and stored at −80 °C before use. The crude membrane (250 µg per sample) was incubated with the inhibitor in an assay buffer (100 mM NaCl and 10 mM HEPES/Tris, pH 7.4) or Na^+^-free assay buffer (100 mM choline chloride and 10 mM HEPES/Tris, pH 7.4) at 25 °C for 4 h. The reactions were terminated by filtering through a GF/C filter plate (Corning) presoaked in assay buffer containing 0.1% bovine serum albumin. The sample in the filter plate was washed thrice with assay buffer and eluted with acetonitrile:water (80:20, v/v). The solution extracted from the filter plate was diluted with formic acid:water (1:1000, v/v) containing verapamil as the internal standard. Inhibitor concentration was quantified using LC–MS/MS.

Nonspecific binding was measured using the crude membrane of non-transfected FreeStyle 293-F cells. Specific binding was calculated by subtracting nonspecific binding from the binding of hB^0^AT1-expressing cells. The equilibrium dissociation constant (*K*_d_) and maximum number of binding sites (*B*_max_) were calculated via nonlinear regression using GraphPad Prism 8.4.3.

### Quantification of B^0^AT1 inhibitors bound to the crude membrane and d5-phenylalanine in cell lysates using LC–MS/MS

The concentrations of the extract solution from the filter plate and cell lysate were quantified using a tandem mass spectrometer QTRAP6500 System (SCIEX) coupled with an ACQUITY UPLC system (Waters) using the internal standard method. Mobile phases A and B consisted of formic acid:water (1:1000, v/v) and acetonitrile, respectively. Chromatographic separation was performed on an ACQUITY UPLC BEH C18 column (2.1 mm × 100 mm, 1.7 μm; Waters) at 50 °C with the following gradient of mobile phase B: 2% (0.00–0.50 min), 2–98% (0.50–2.00 min), 98% (2.00–2.50 min), and 2% (2.51–3.00 min); the flow rate was 0.4 mL/min. Mass spectrometric detection was performed by multiple reaction monitoring in the electrospray ionization positive-ion mode controlled by Analyst 1.6.2, using m/z 255.3/102.8 for compound 1; 301.2/199.0 for compound 2; 356.2/198.9 for compound 3; 171.2/125.0 for d5-phenylalanine; and 174.2/128.0 for d8-phenylalanine.

### Preparation of compound and phenylalanine administration solution

The test compound was weighed and suspended in 0.5% carboxymethyl cellulose. L-phenylalanine (Sigma-Aldrich) was dissolved in physiological saline to prepare a 25 mg/mL solution. This solution was prepared on the day of phenylalanine administration.

### Administration of compound and collection of biospecimen

Male *Pah^enu2^* mice (12 weeks old, The Jackson Laboratory Japan, Inc., Yokohama, Japan) were used as test animals. Two days before the experiment, mice were placed in metabolic cages for urine collection. After habituation, mice were randomly assigned into two groups with matched mean body weights and variances, verified by an in-house algorithm. The test compound was orally administered at a dose of 30 mg/10 mL/kg. At 4 h after administration, urine samples were collected, and blood samples were collected from the tail vein. Blood was collected in a heparinized capillary tube and centrifuged to obtain plasma.

### Quantification of phenylalanine in mouse urine and plasma via LC–MS/MS

Mouse plasma and urine concentrations were quantified using a tandem mass spectrometer (Quattro Premier XE, Waters, Milford, Massachusetts, USA) coupled to an ACQUITY UPLC system (Waters, Milford, MA, USA) using the internal standard method. Samples were diluted in saline, deproteinized with acetonitrile containing d5-phenylalanine as an internal standard, and centrifuged, and the supernatant was subjected to LC–MS/MS. Mobile phases A and B were heptafluorobutyric acid:water (0.25/1000, v/v) and acetonitrile, respectively. Chromatographic separation was performed on an ACQUITY UPLC HSS C18 column (2.1 mm × 100 mm, 1.8 µm; Waters) at 50 °C, with the following gradient of mobile phase B: 2% (0.00–0.50 min), 2–98% (0.50–3.50 min), 98% (3.50–4.25 min), and 2% (4.26–5.00 min); the flow rate was 0.4 mL/min. Mass spectrometric detection was performed using multiple reaction monitoring in the electrospray ionization positive-ion mode controlled by MassLynx using m/z 166.1/120.0 for phenylalanine and 171.2/125.0 for d5-phenylalanine.

### Statistical analysis

For the *in vivo* experiments, statistical analyses were performed using SAS 10.1 (SAS Institute, Inc.). The effect of test compounds on Phe excretion in the urine and levels in the plasma was analyzed using an unpaired, two-tailed Student’s *t*-test for comparisons between two groups. p-values < 0.01 were considered statistically significant.

## Supporting information

Supplymentaly Fig.1-8, Supplementary Data Table 1-3, Supplementary Protocols

## ACKNOWLEDGMENTS

We thank the scientific staff at the cryo-EM facility of the University of Tokyo, as ell as all members who contributed to this project. We also thank Sohyaku Innovative Research Division, Mitsubishi Tanabe Pharma Co., Ltd for providing facility support. The authors received no specific funding for this study.

## AUTHOR CONTRIBUTIONS

T.I., H.T., and W.S. designed the compounds and performed the structure-activity relationship analyses. Y.I. designed and performed the *in vivo* model evaluation analyses; M.H. and I.M. performed the cryo-EM analyses with sample preparation assistance from T.K. and H.M. T.A. designed and performed the functional analyses with sample preparation assistance from T.K. H.M., and I.M. performed model building and refinement. T.I., T.A., I.M., M.H., and O.N. supervised the study and wrote and edited the manuscript with help from all other authors. T.I., T.A., I.M., M.H., and O.N. supervised the study.

## MATERIALS & CORRESPONDENCE

### DATA AVAILABILITY

Cryo-EM density maps were deposited in the Electron Microscopy Data Bank under the accession codes EMD-62632 (B^0^AT1—ACE2 complex with compound 1), EMD-62627 (B^0^AT1 with compound 1 focused refinement), EMD-62633 (B^0^AT1–ACE2 complex with compound 2), EMD-62628 (B^0^AT1 with compound 2 focused refinement), EMD-62634 (B0AT1–ACE2 complex with compound 3), EMD-62629 (B^0^AT1 with compound 3 focused refinement), EMD-62635 (apo B^0^AT1–ACE2 outward-facing), EMD-62630 (apo B^0^AT1 outward-facing focused refinement), EMD-63360 (B^0^AT1 antibody-bound inward-open conformation) and EMD-62631 (apo B^0^AT1 inward-open conformation focused refinement). The atomic coordinates have been deposited in the Protein Data Bank under IDs 9KXY (B^0^AT1–ACE2 complex with compound 1), 9KXT (B^0^AT1 with compound 1 focused refinement), 9KXZ (B^0^AT1–ACE2 complex with compound 2), 9KXU (B^0^AT1 with compound 2 focused refinement), 9KX0 (B^0^AT1–ACE2 complex with compound 3), 9KXV (B^0^AT1 with compound 3 focused refinement), 9KY1 (apo B^0^AT1–ACE2 outward-facing), 9KXW (apo-B^0^AT1 outward-facing focused refinement), 9LSZ (B^0^AT1 antibody-bound inward-open conformation) and 9KXX (apo B^0^AT1 inward-open conformation focused refinement). The source data are provided in this study.

---

**Extended Data Fig. 1.**
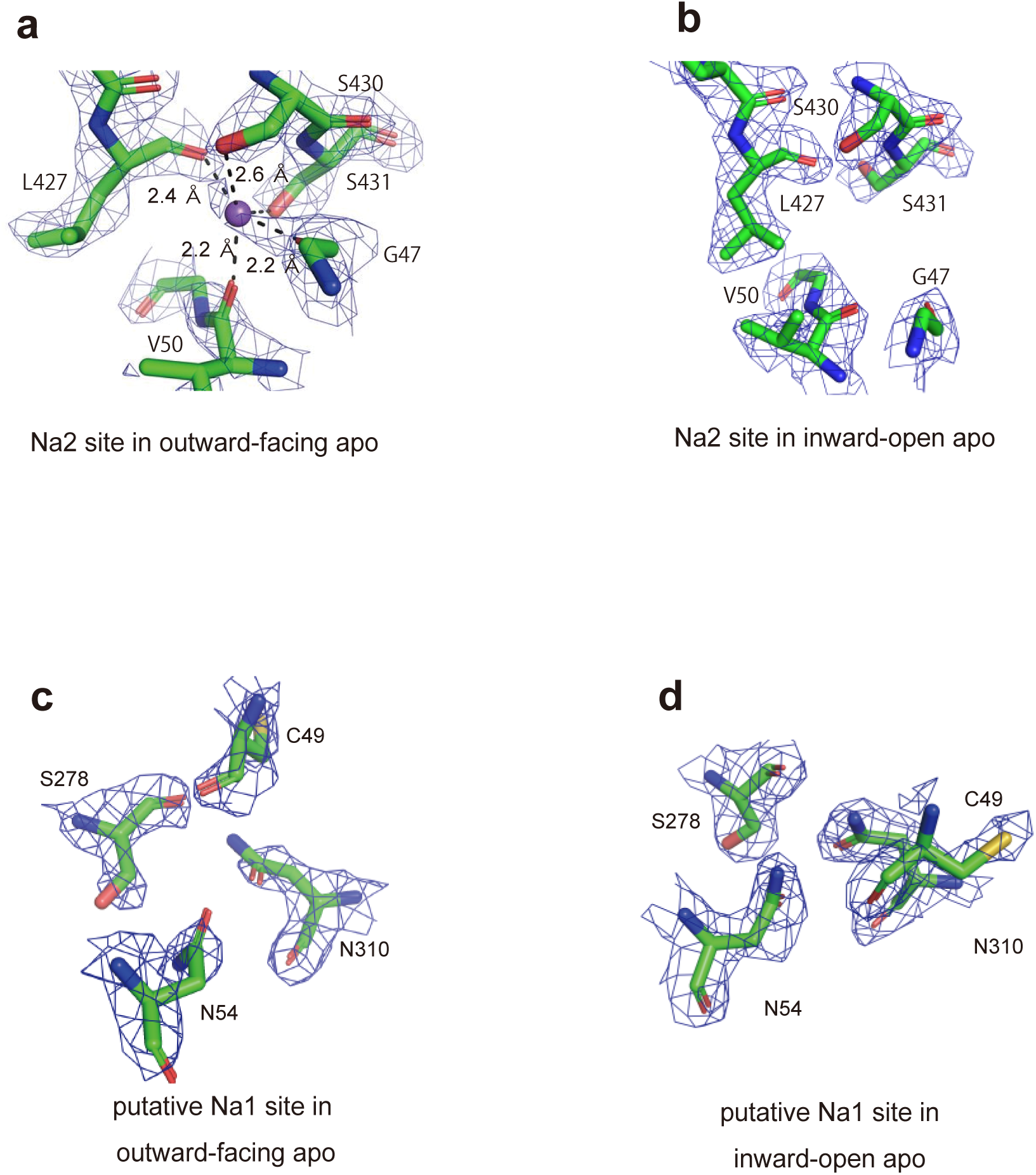
Cryo-EM density maps of apo Na2 site, and apo putative Na1 site in B⁰AT1. a) Na2 site of the outward-facing apo B^0^AT1, with five surrounding atoms coordinated to Na^+^. b) Inward-open form of apo B^0^AT1. c, d) The putative Na1 sites of the the outward-facing and inward open apo B^0^AT1 show no density map corresponding to Na^+^. Contour levels of cryo-EM density are 2.5σ(a), and 2.0σ(b-d).

**Extended Data Fig. 2.**
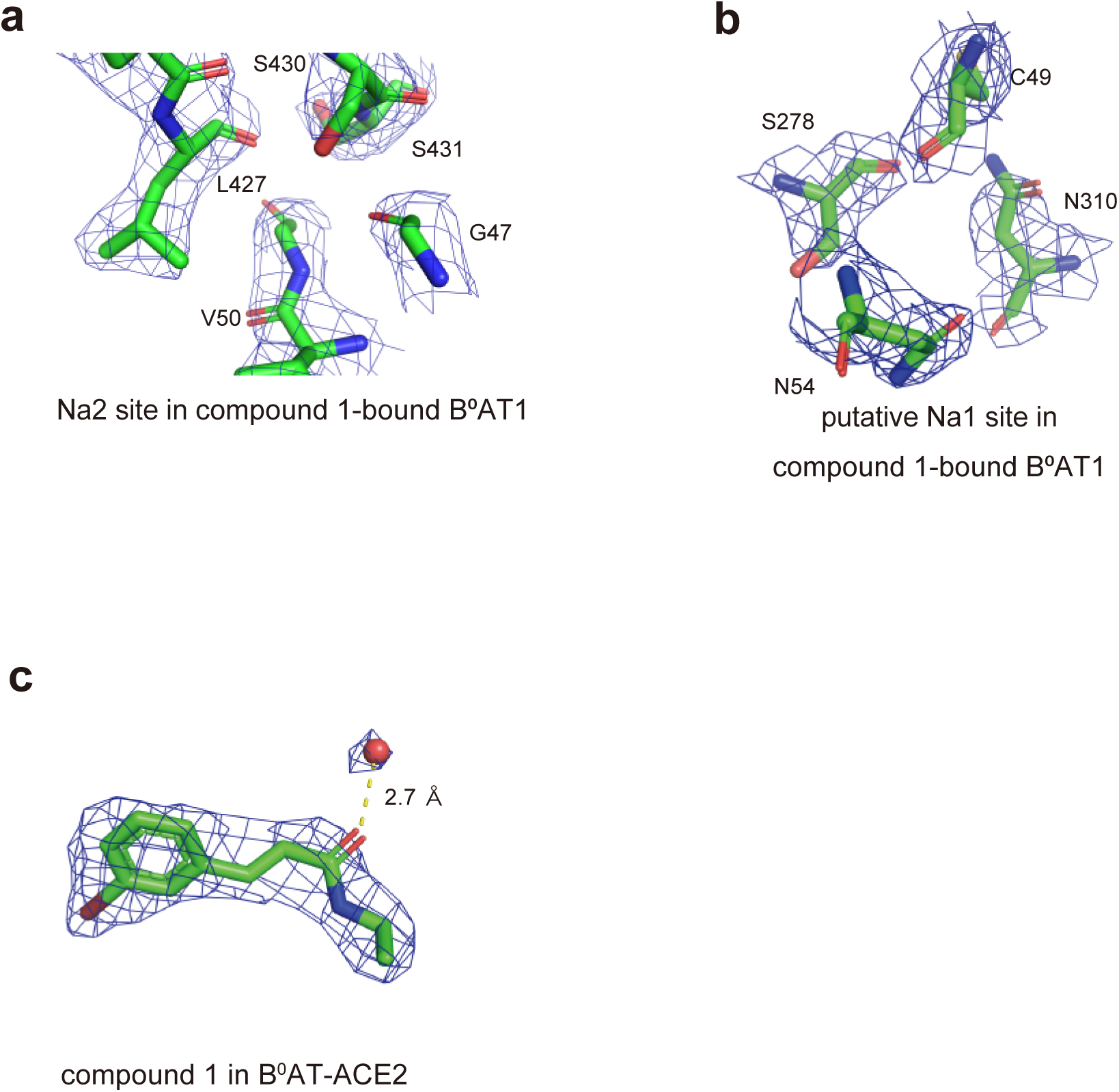
Cryo-EM density maps of Na2 and putative Na1 sites in compound 1-bound B⁰AT1. a) The Na2 site, b) The putative Na1 site of the compound 1-bound structure. d) Compounds 1 and its bound water molecules. Contour levels of cryo-EM density are 2.0σ(d).

**Extended Data Fig. 3.**
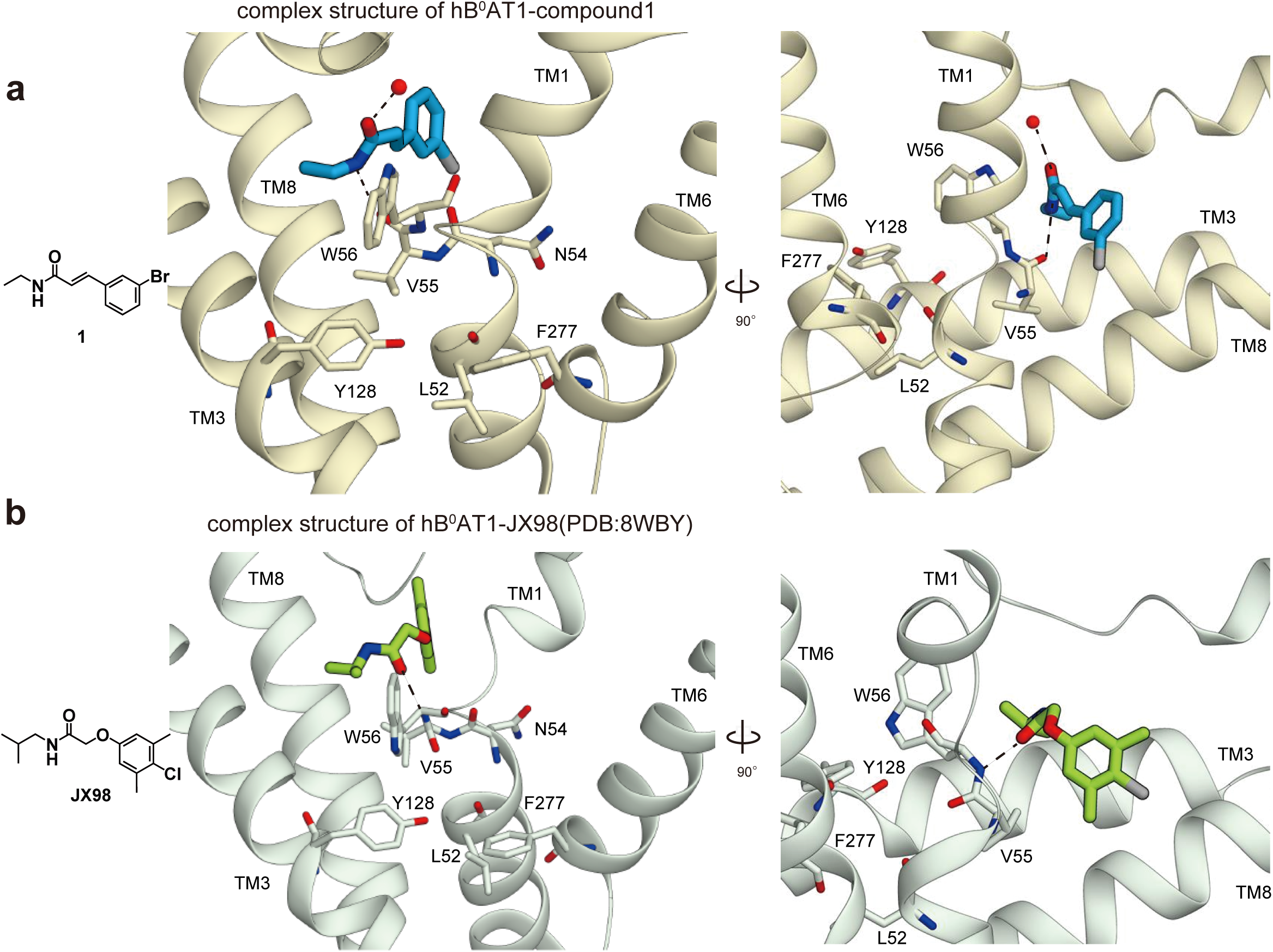
Comparison of binding sites for compound 1, and JX98. a) The amide group of compound 1 forms a hydrogen bond with V55. b) The carbonyl of JX98 directs to W56 to form a hydrogen bond. The complex structure of hB⁰AT1-JX98 was generated using PDB:8WBY

**Extended Data Fig. 4.**
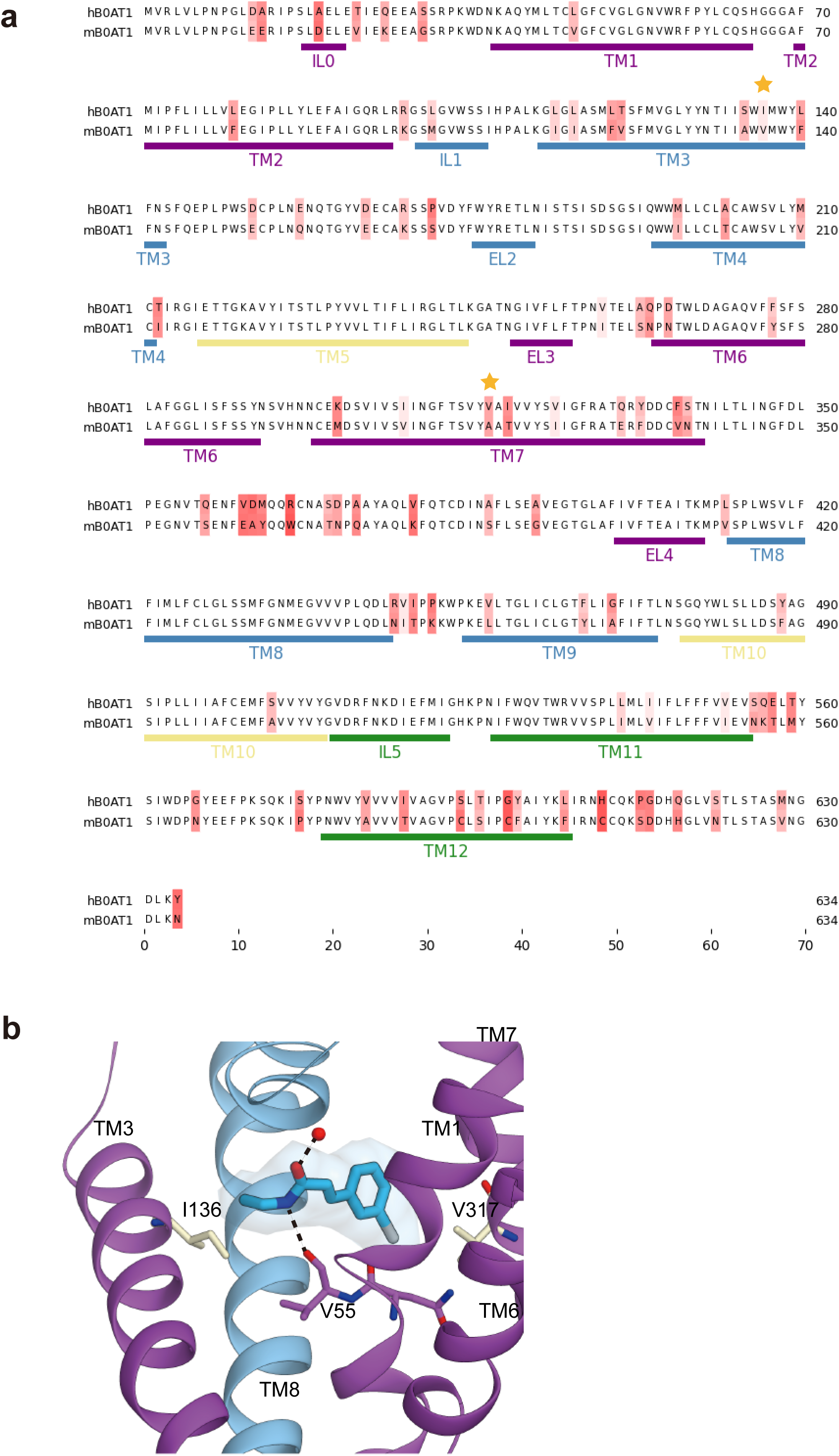
Amino acid sequence comparison between human and mouse. a) Amino acid sequence comparison between human and mouse. The star represents the amino acids that differ between hB⁰AT1 and mB⁰AT1 around the compound 1 binding site. b) Complex structure of hB^0^AT1-compound1. Amino acids around the binding site of compound 1 that differ from mouse are indicated in white.

**Extended Data Fig. 5.**
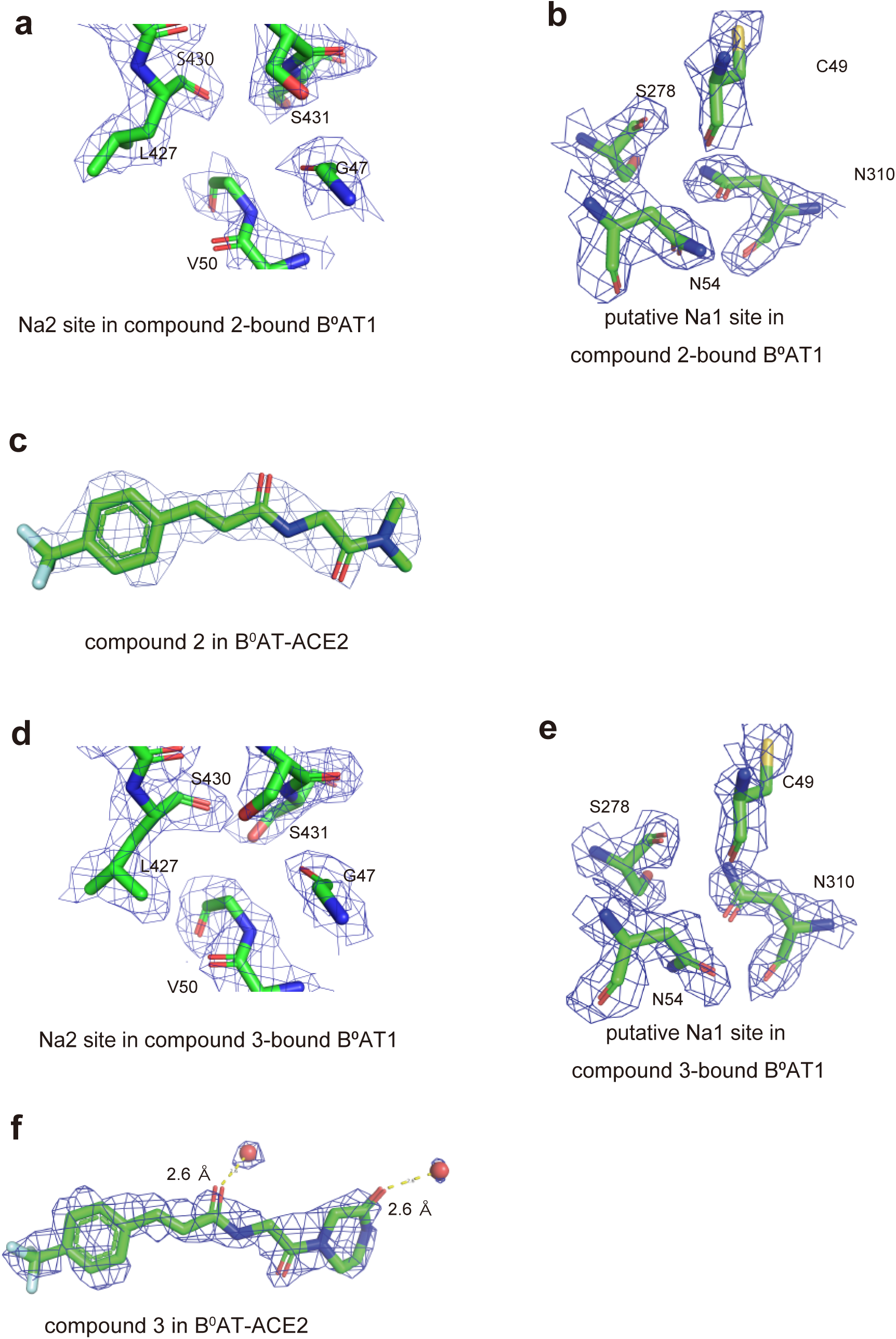
Cryo-EM density maps of Na2, putative Na1 sites and compounds in compound 2, 3-bound B⁰AT1 structures. a) The Na2 site, b) The putative Na1 sites in compound 2-bound B⁰AT1 and c) Compounds 2 and its bound water molecules. d) The Na2 site, e) The putative Na1 sites in compound 3-bound B⁰AT1 and f) Compounds 3 and its bound water molecules. Contour levels of cryo-EM density are 2.0σ.

**Extended Data Fig. 6.**
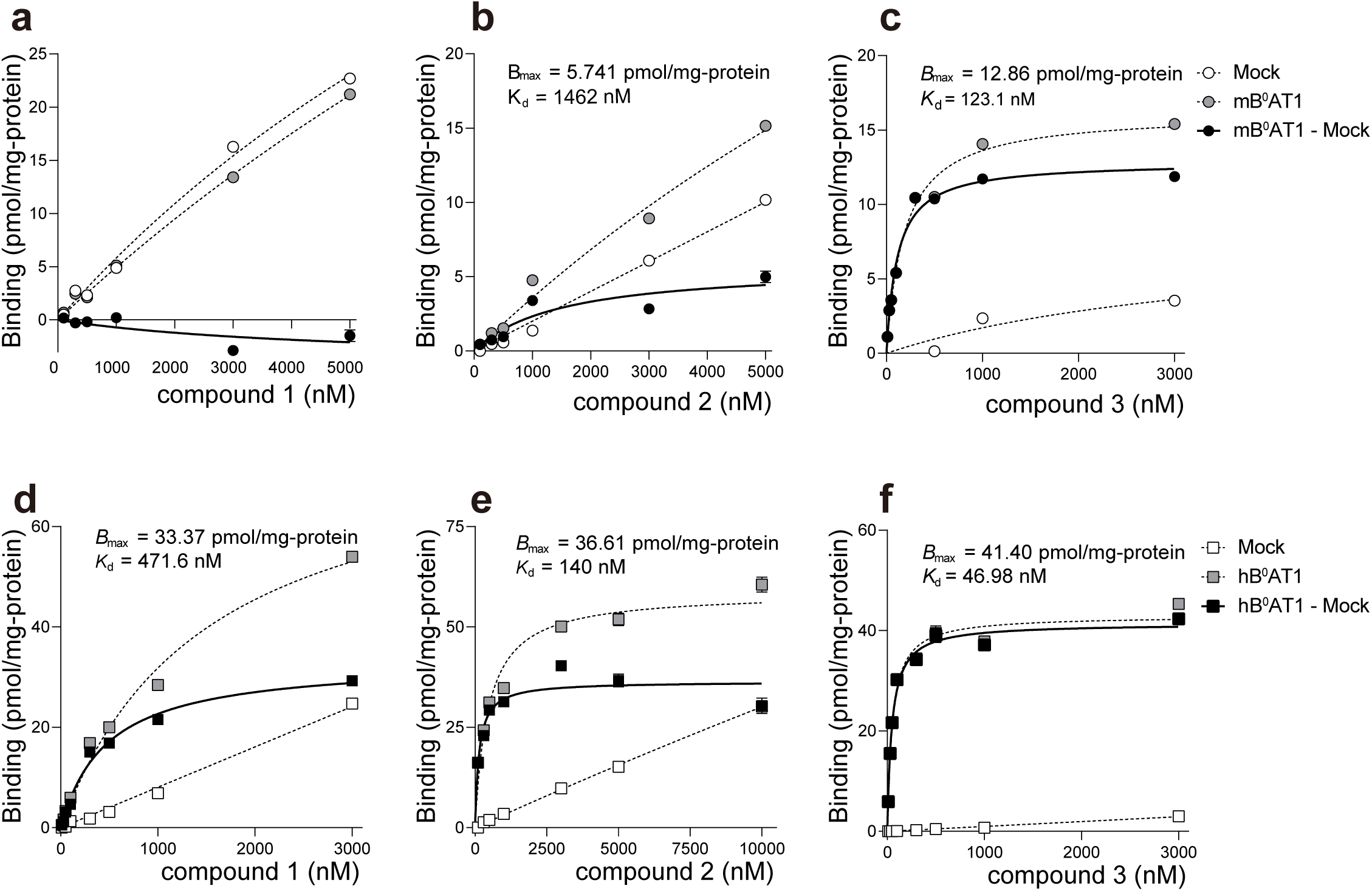
Comparison of binding affinity of compound 1-3 to human and mouse B⁰AT1. a-c) Binding affinity of compound 1-3 using Mock cell and mB⁰AT1 and ACE2-expressing cell. d-f) Binding affinity of compound 1-3 using Mock cell and hB⁰AT1 and ACE2-expressing cell. Specific binding (black) was calculated by subtracting the binding of mock cells (white) from the binding of B⁰AT1 and ACE2 expressing cells (gray). Each point represents the mean ± s.e.m. (n = 3, technical replicates).

**Extended Data Fig. 7.**
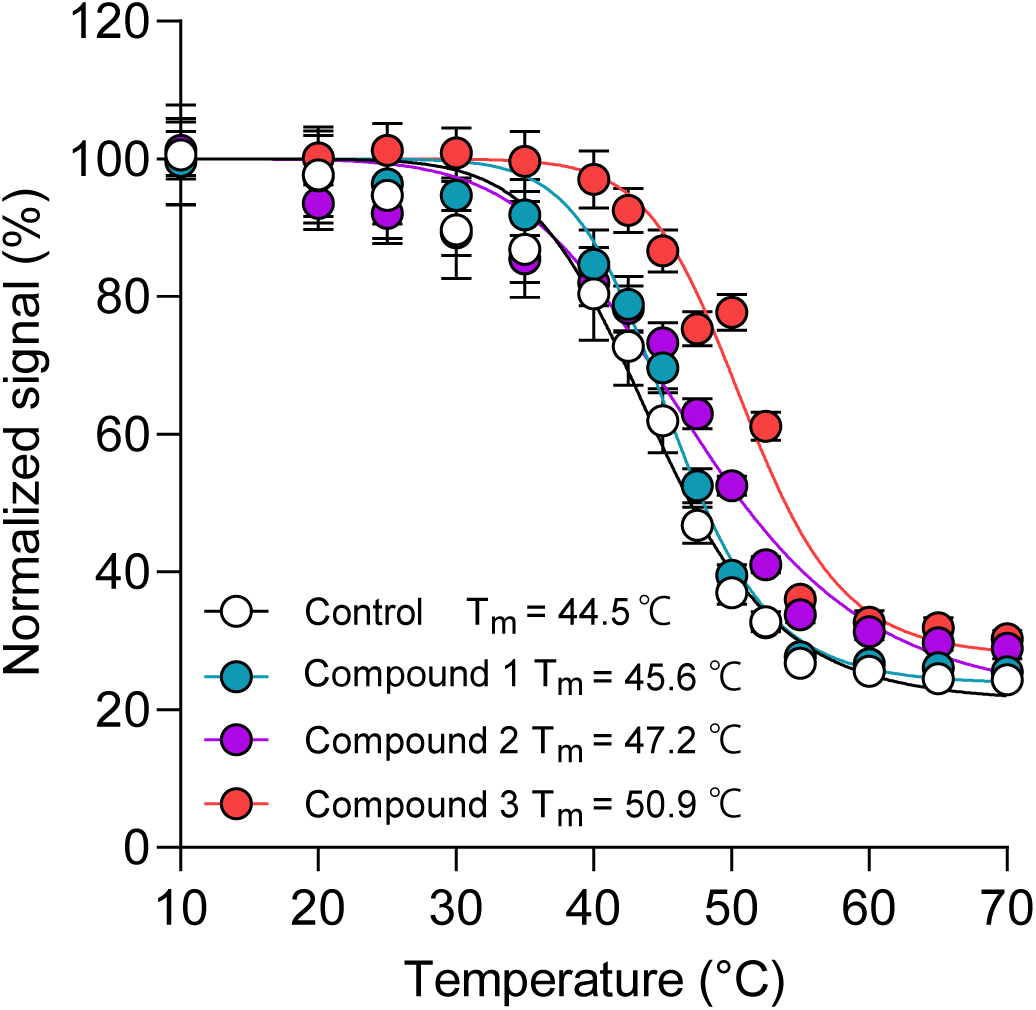
Confirmation of thermal stabilization effect using FSEC. FSEC-based thermal shift assay using purified hB^0^AT1-ACE2 in the presence or absence of inhibitors. Each point represents the mean ± s.e.m. (n = 3, technical replicates).

**Extended Data Fig. 8.**
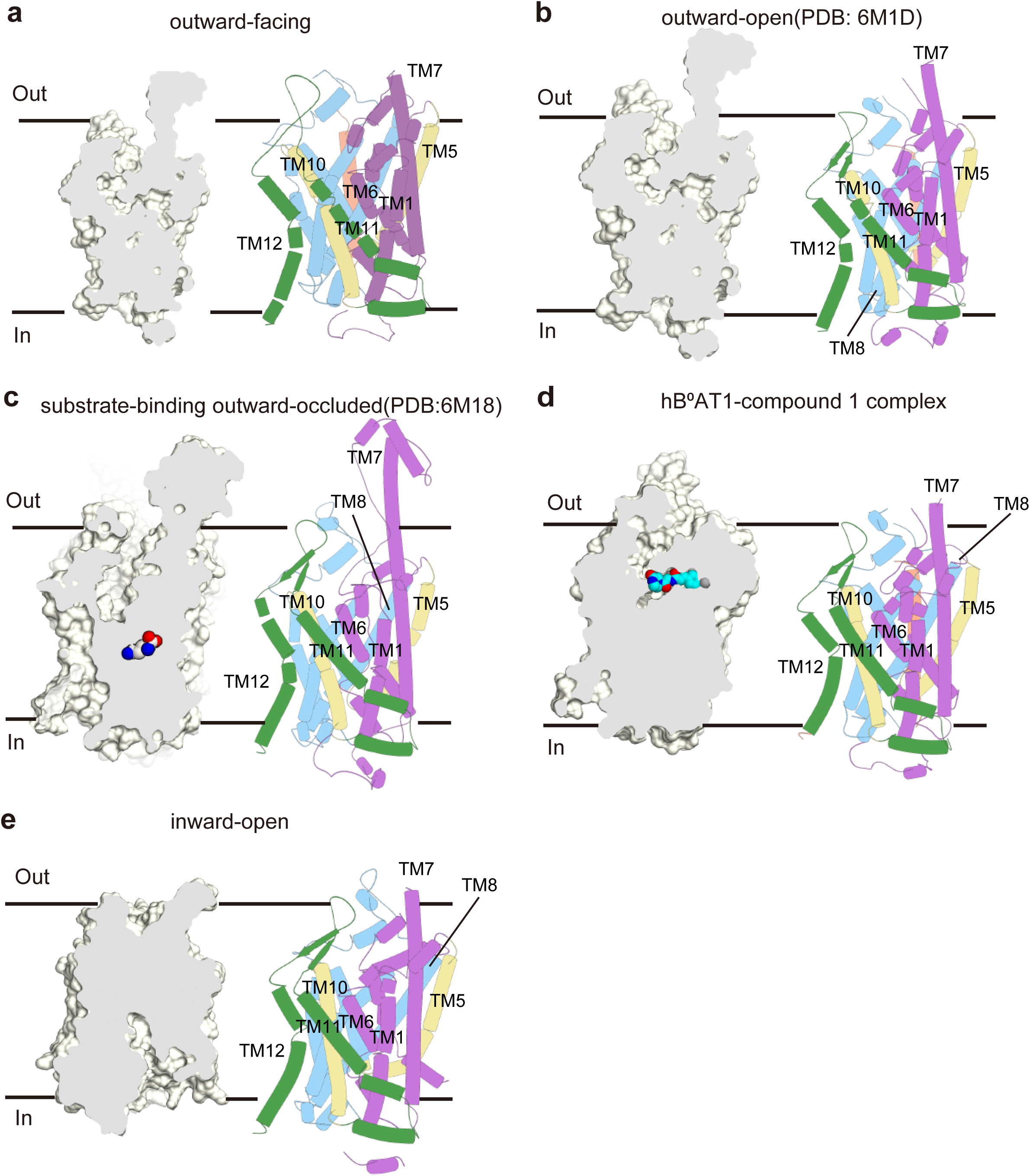
Comparing structure of state for hB⁰AT1. a) Outward-facing b) outward open. c) substrate-binding outward-occluded, d) compound1-bound outward-facing, e) antibody-bound inward-open structures. For the cylindal structures, the bundle domain (TM1,2,6,7) is in purple, the hash domain (TM3,4,8,9) in blue, the gate helices (TM5,10) in yellow, the rest of the protein (TM11, 12) in green. The cavity for compound binding appears only in an outward-open conformation when there is no compound.

**Extended Data Fig. 9.**
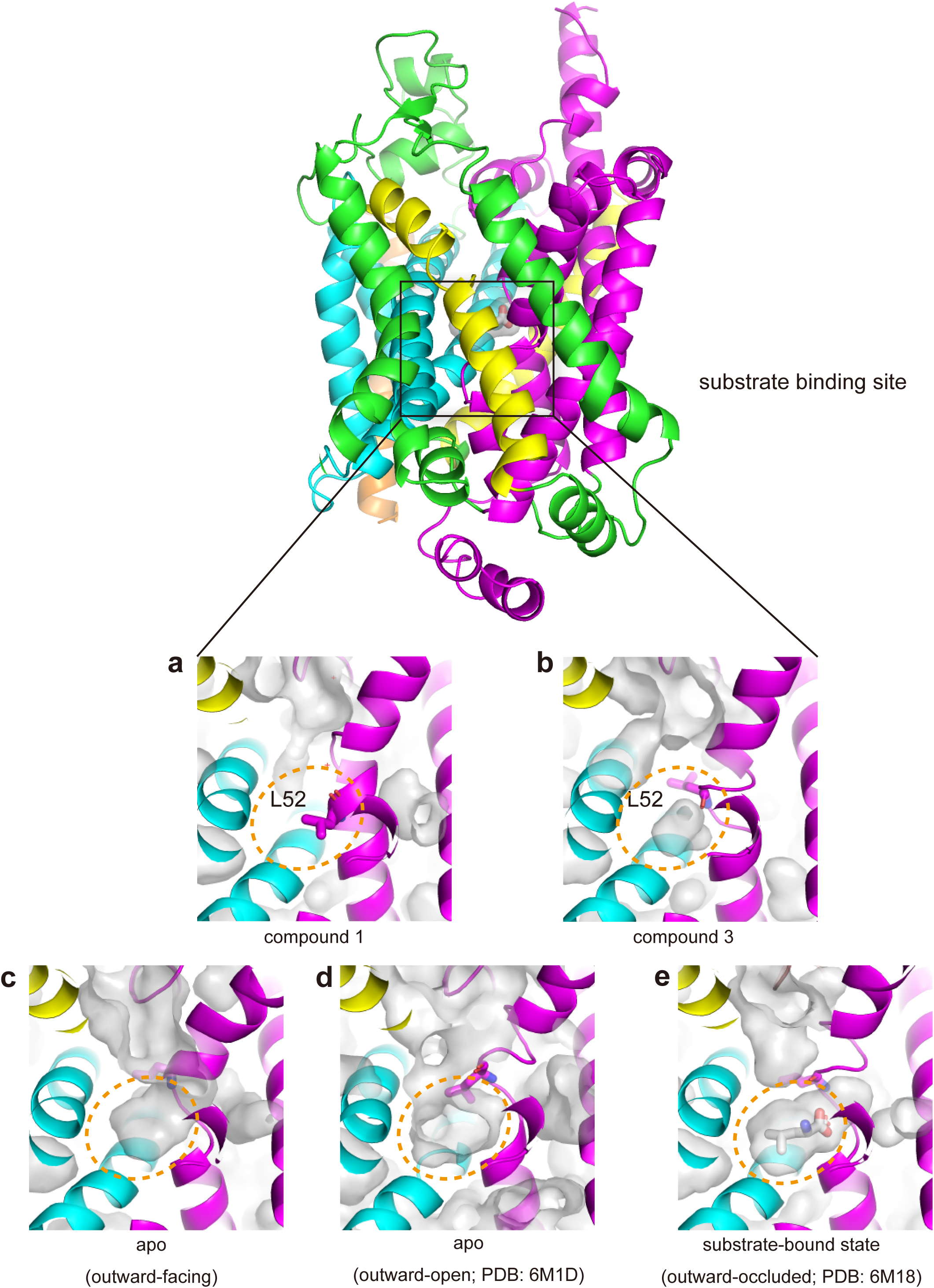
Structure of substrate binding sites in each state. a) When compound 1 binds, the side chain of L52 flips, and the substrate binding site is completely lost. b) In the case of compound 3, the side chain of L52 does not flip, so an occluded space remains. c, d) The outward-facing, outward-open structure indicates that the substrate binding site is connected to the outside of the cell. e) The substrate binding state indicates that the substrate is in the occluded space.

**Extended Data Fig. 10.**
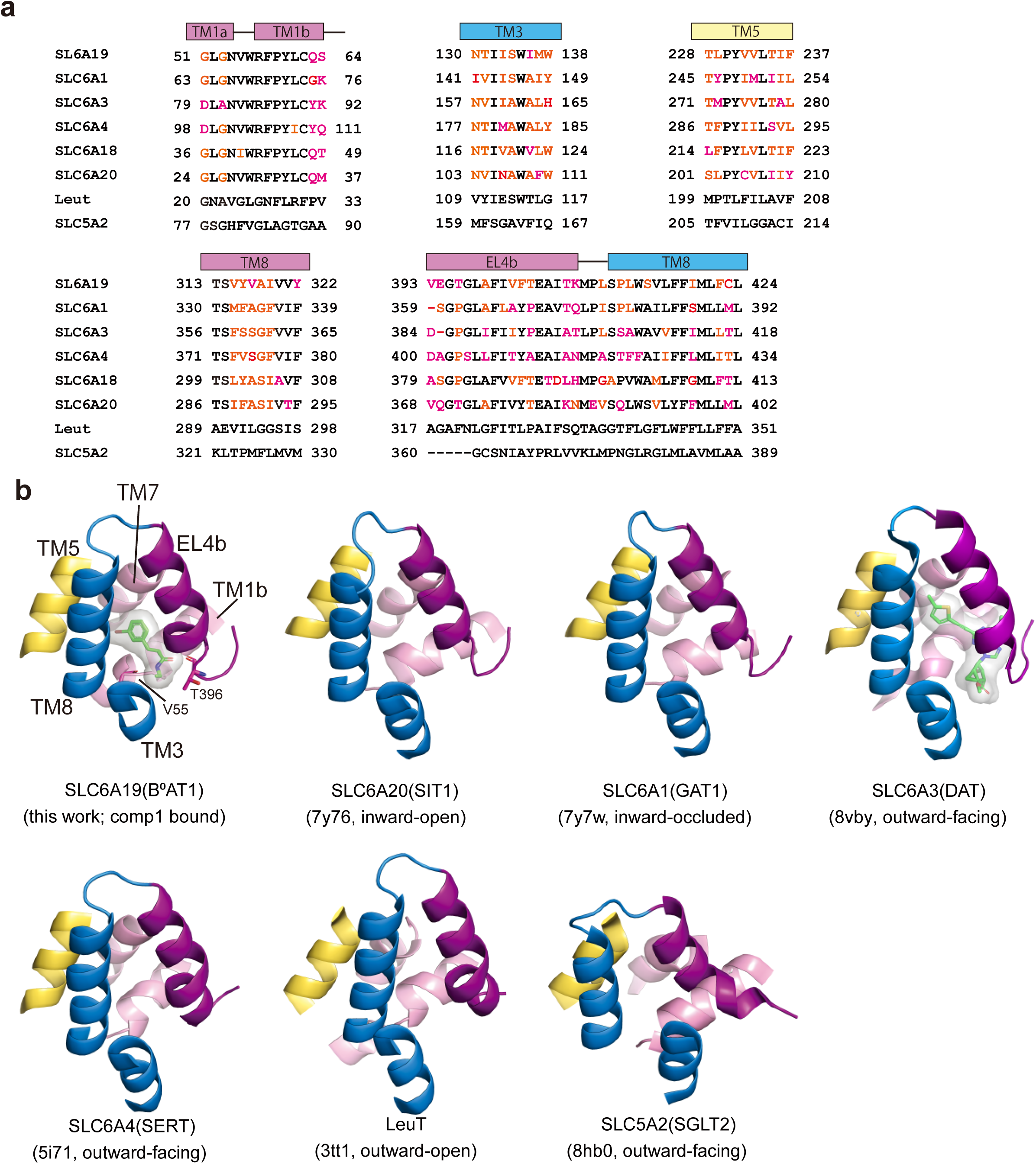
Comparison of the allosteric sites among SLC5 and SLC6 transporters. a) Sequence comparison: The six α-helices that make up the site are conserved in all the transporters. b) Structural comparison: The binding site formed by the six α-helices is conserved in all transporters. Other than B⁰AT1, it has been reported that a compound is bounded to the same site of DAT(SLC6A3).

