## Supplementary material for "Structure-guided development of a potent human B^0^AT1 inhibitor effective in a mouse model of phenylketonuria": Supplymentaly Fig.1-8, Supplementary Data Table 1-3, Supplementary Protocols

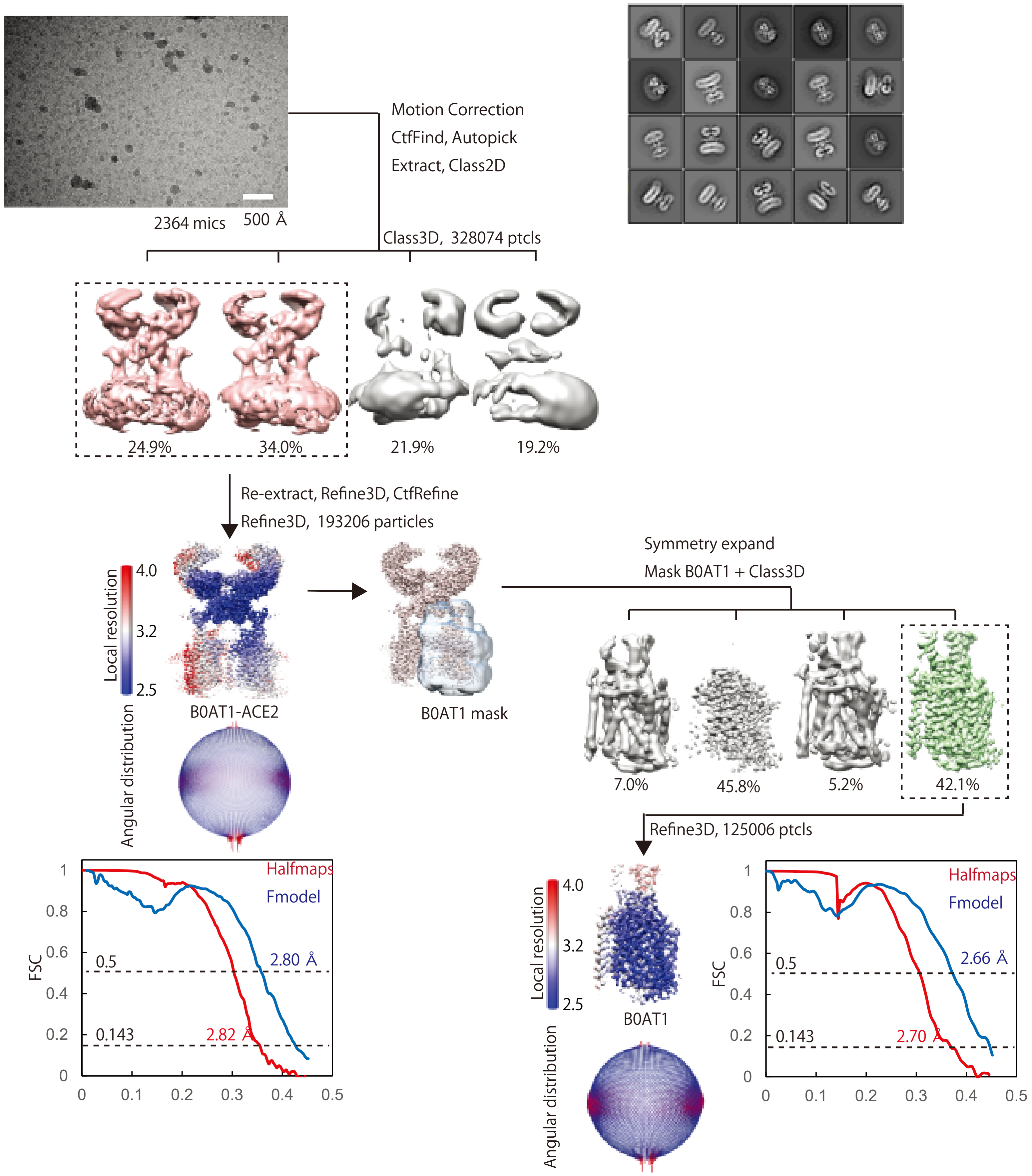


**Supplementary Figure 1** Structure determination of the compound 1 bound B⁰AT1-ACE2

Flow chart for determinaing the structure of the B⁰AT1-ACE2 complex in the presence of compound 1.

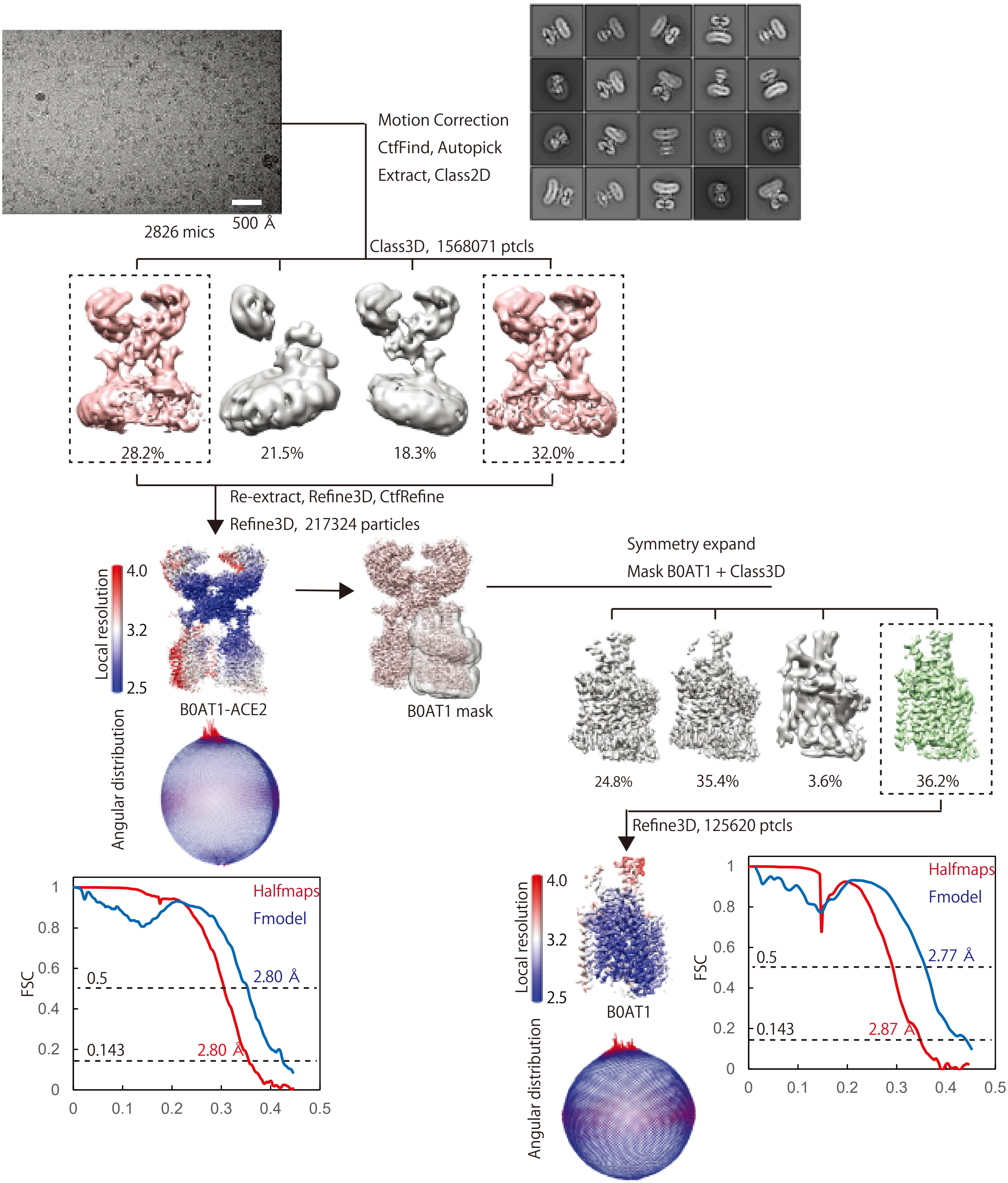


**Supplementary Figure 2** Structure determination of the compound 2 bound B⁰AT1-ACE2

Flow chart for determinaing the structure of the B⁰AT1-ACE2 complex in the presence of compound 2.

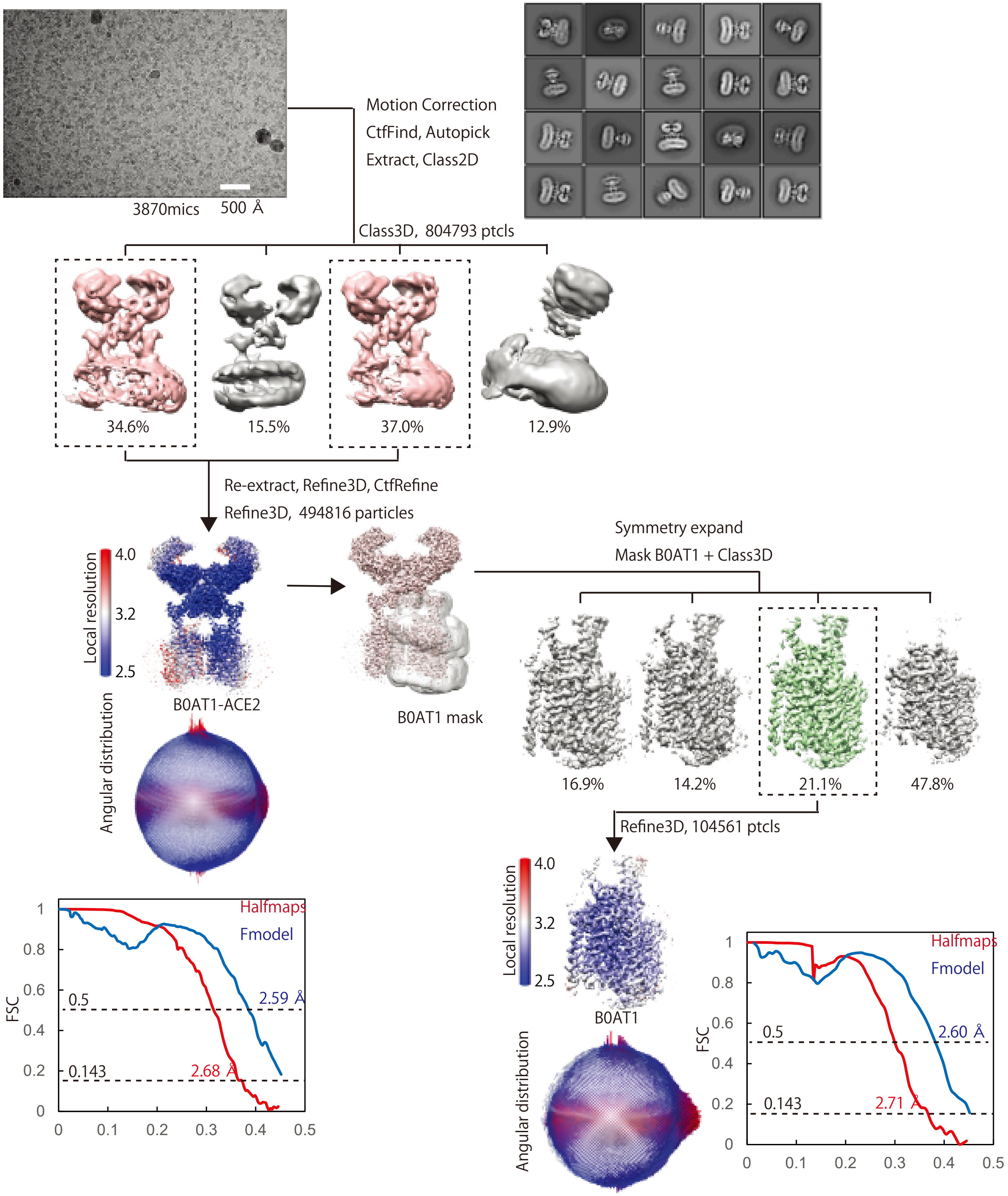


**Supplementary Figure 3** Structure determination of the compound 3 bound B⁰AT1-ACE2

Flow chart for determinaing the structure of the B⁰AT1-ACE2 complex in the presence of compound 3.

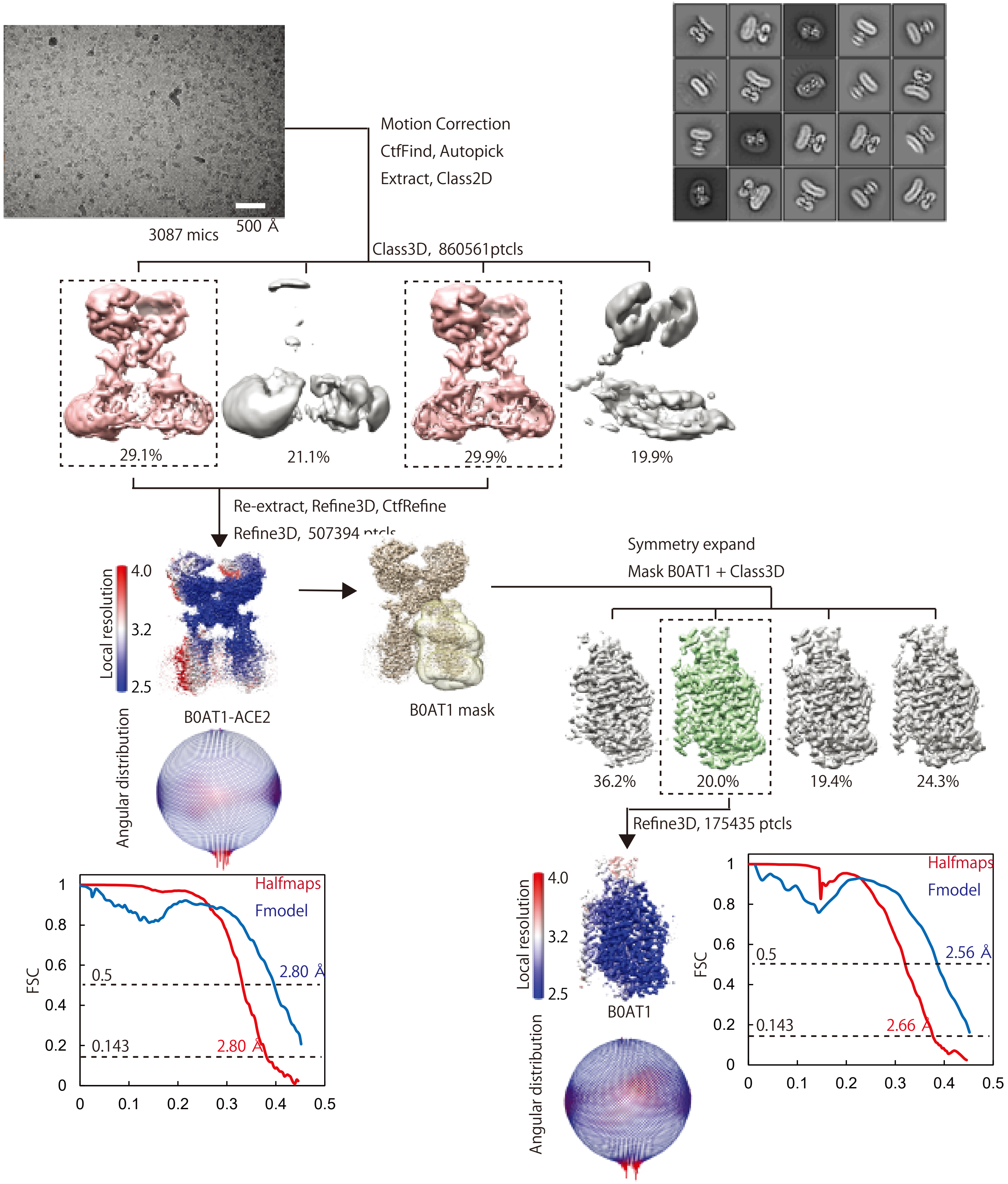


**Supplementary Figure 4** Structure determination of apo BºAT1-ACE2

Flow chart for determinaing the structure of the BºAT1-ACE2 complex.

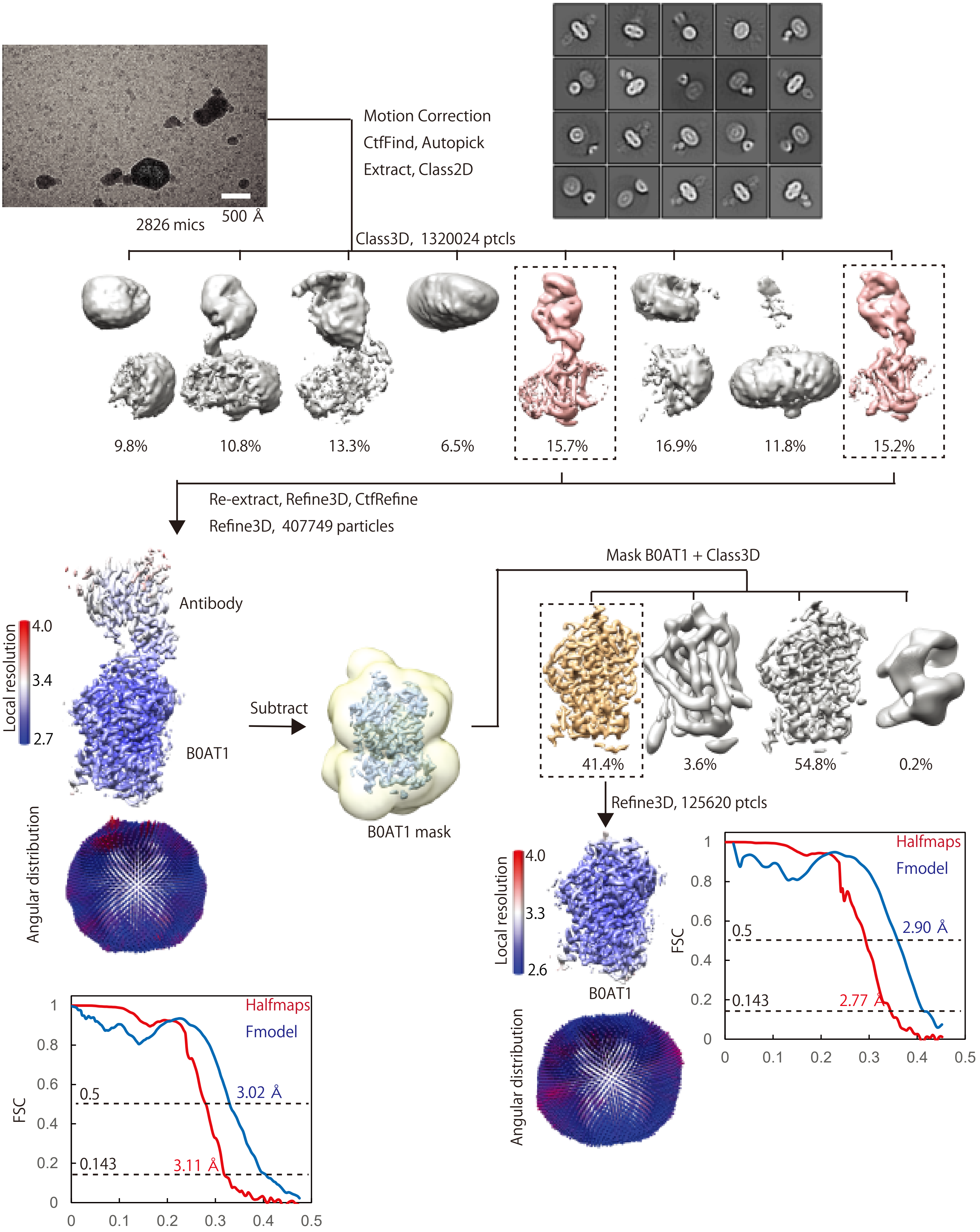


**Supplementary Figure 5** Structure determination of the inward-open form of apo B⁰AT1

Flow chart for determinaing the structure of the inward-open form of apo B⁰AT1 complexed with antibody.

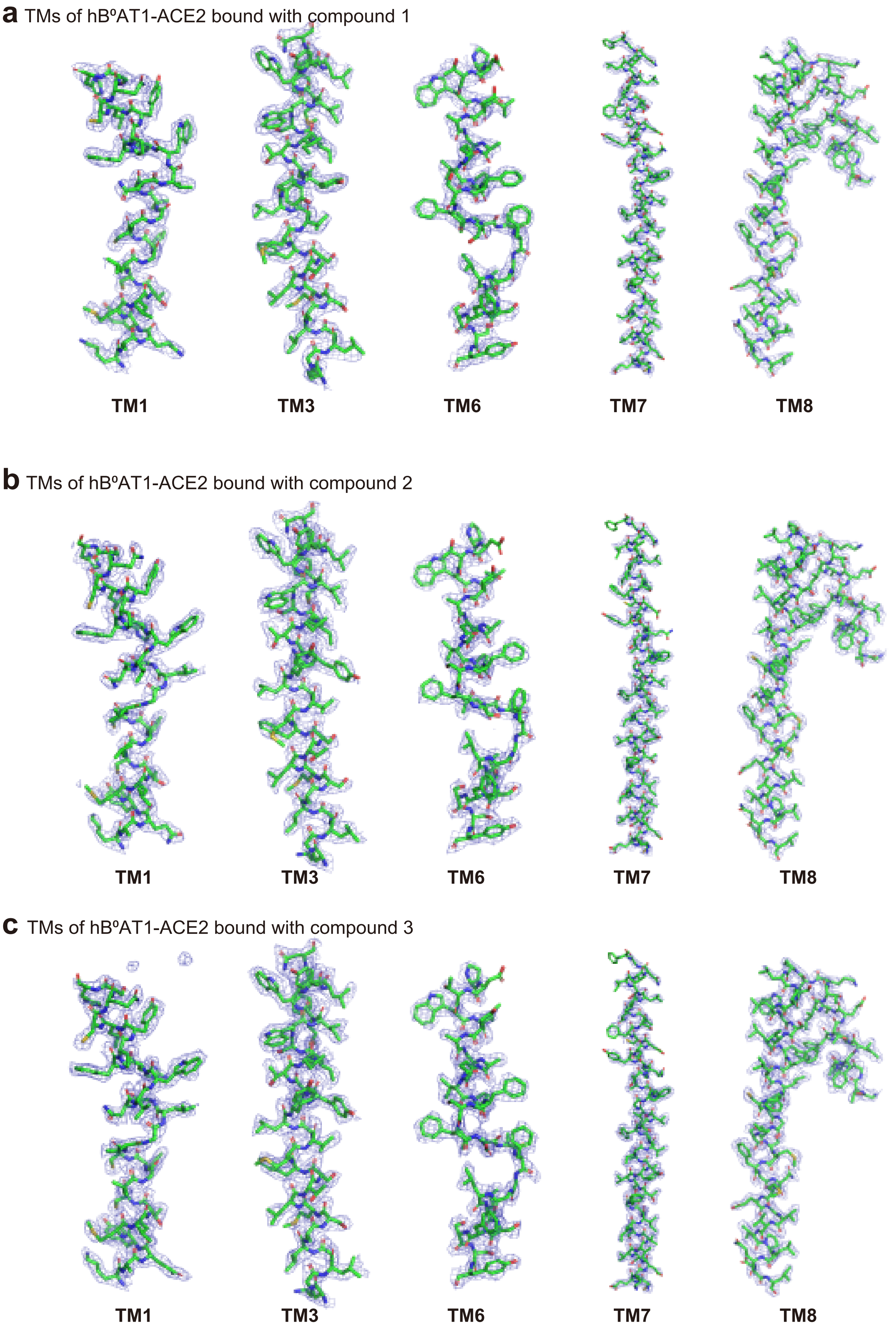


**Supplementary Figure 6** Cryo-EM density maps of ACE2-hB$^0$AT1 bound with inhibitors

Cryo EM density maps shown at threshhold of 2s(a-c).

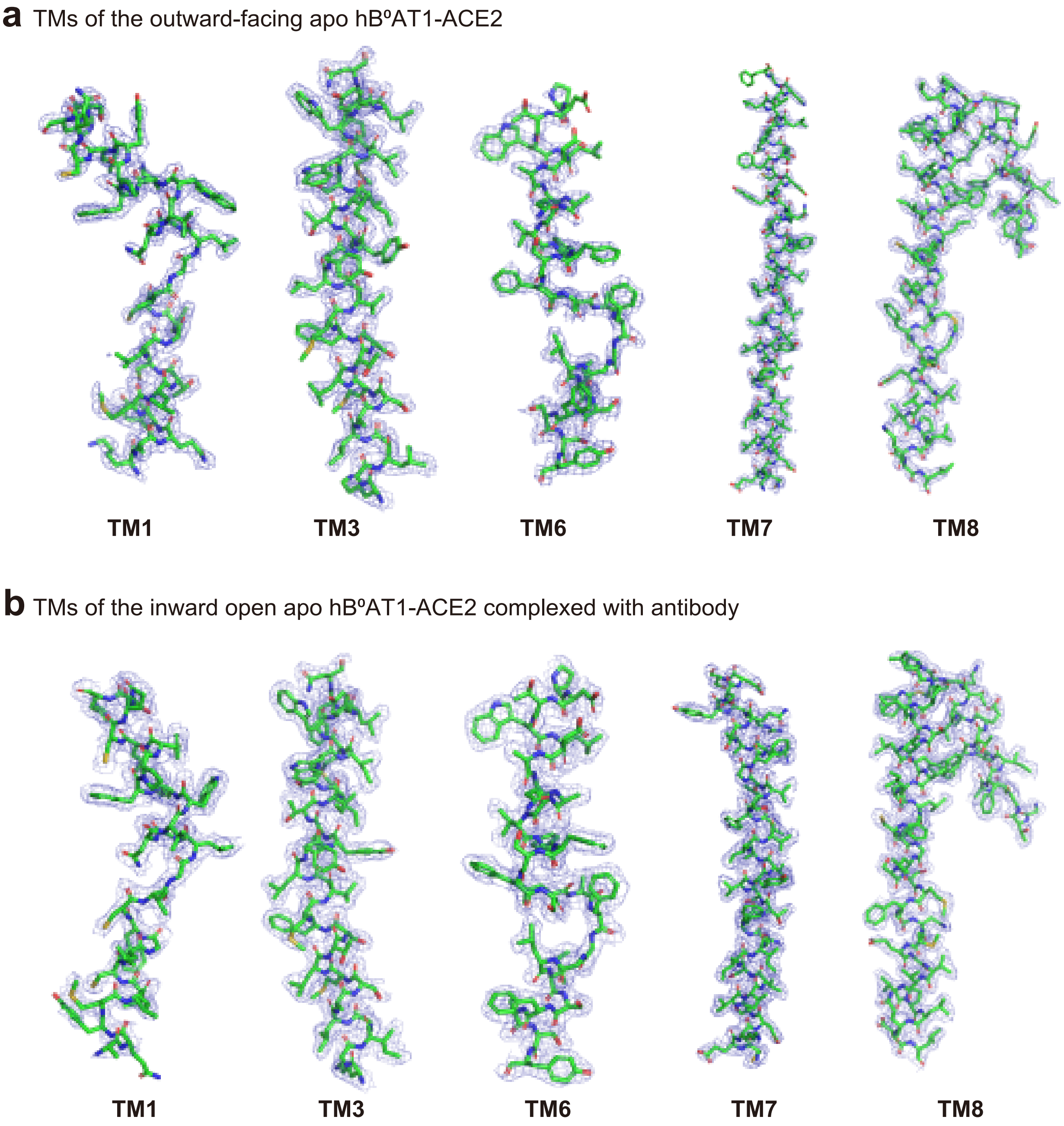


**Supplementary Figure 7** Cryo-EM density maps of apo hB$^0$AT1

Contour levels of cryo-EM density are 1.0s(a-b).

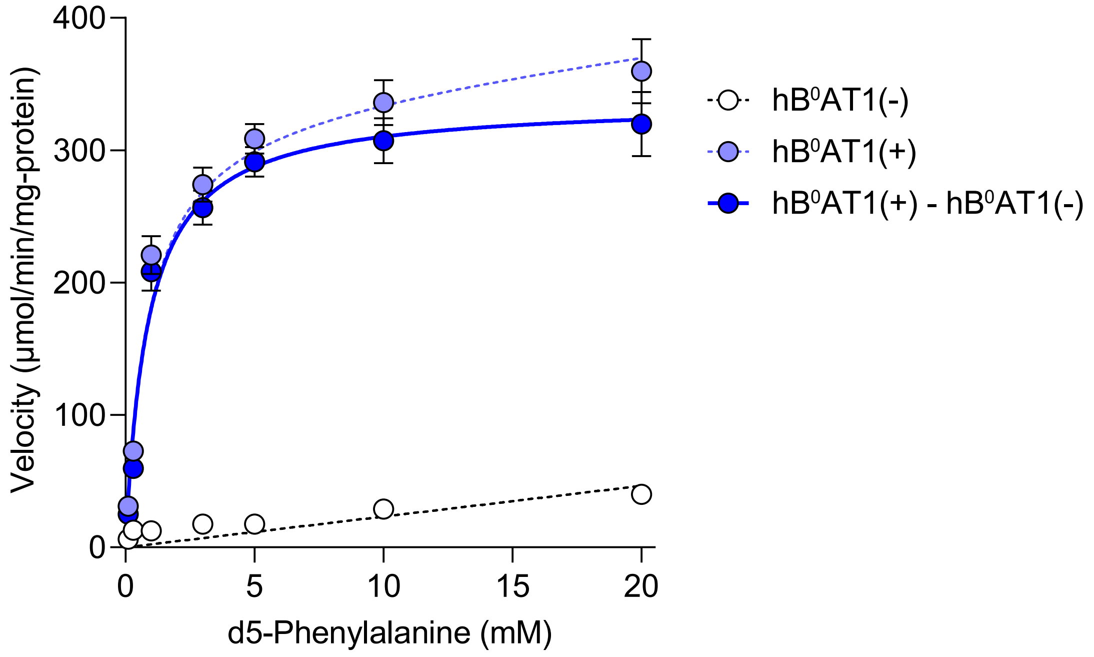


**Supplementary Figure 8** Concentration dependence of d5-phenylalanine uptake by hBºAT1 and ACE2-expressing cells and non-expressing cells.

BºAT1-mediated uptake was calculated by subtracting the uptake by non-expressing cells from that of hBºAT1 and ACE2-expressing cells. Each point represents the mean ± s.e.m. (n = 3, biological replicates).

| | compound 3 | 0 nM | 1500 nM | 4000 nM |
|---|---|---|---|---|
| d5-Phenylalanine | $V_{max}$ (μmol/min/mg-protein) | 337 ± 12 | 351 ± 26 | 380 ± 37 |
| | $K_m$ (mM) | 0.866 ± 0.144 | 5.19 ± 1.01 | 12.1 ± 2.3 |

**Supplementary Data Table 1** Kinetic parameters of d5-phenylalanine uptake by hB$^{o}$AT1 in the presense of compound 3.
Each data represents the mean ± s.e.m. (n = 3, technical replicates).

| | | Na(+) | Na(-) |
|---|---|---|---|
| compound 3 | $B_{max}$ (pmol/mg-protein) | 634 ± 18 | 462 ± 13 |
| | $K_d$ (nM) | 38.0 ± 5.2 | 21.9 ± 3.4 |

**Supplementary Data Table 2** Kinetic parameters of compound 3 in the crude membrane expressing B$^o$AT1.
Each data represents the mean ± s.e.m. (n = 3, technical replicates).

| Time (h) | Plasma concentration (ng/mL) |
|---|---|
| 4 | 915 ± 311 |
| 8 | 1590 ± 470 |

**Supplementary Data Table 3** Plasma concentration of compound 3 in *Pah*$^{enu2}$ mice

Plasma concentration of compound 3 after administration orally at a dose of 30 mg/kg in *Pah*$^{enu2}$ mice. Each data represents the mean ± s.d. (n = 8, biological replicates).

**General Methods for Chemistry.** The progress of the reactions was monitored using analytical thin layer chromatography (TLC) on glass plates (TLC Silica gel 60 F254), and the products were visualized with UV light or LC/MS (Waters, ACQUITY QDa or ACQUITY SQD). For column chromatography, silica gel (Wakogel® 60 N, FUJIFILM Wako, Osaka, Japan) or a pre-packed silica gel column (Hi-Flash-M, L, 2L, 3 L or 5L, YAMAZEN, Osaka, Japan) was used. Nuclear magnetic resonance (NMR) spectra were recorded on a Bruker AVANCE 400 spectrometer (400 MHz) at 400 and 100 MHz for $^{1}H$ and $^{13}C$, respectively, and referenced to tetramethylsilane as an internal standard. NMR data were processed using ACD/Spectrus Processor software and recorded as follows: $^{1}H$ NMR − chemical shift (δ, ppm), multiplicity (s, singlet; d, doublet; t, triplet; q, quartet; m, multiplet), coupling constant (Hz), and integration; $^{13}C$ NMR − chemical shift (δ, ppm). High-resolution mass spectra (HRMS) were recorded using a Thermo Scientific LTQ Orbitrap XL Exact mass spectrometer. The purity of biologically tested compounds was assessed by ultra-high performance liquid chromatography (UHPLC) analysis at 200–400 nm. The UHPLC analysis was performed on a Shimadzu Nexera X2 system with a Waters ACQUITY UPLC BEH C18 (1.7 μm, 2.1 × 100 mm) column.

**Experimental procedure for compound 2, 3.**

**Schemes**

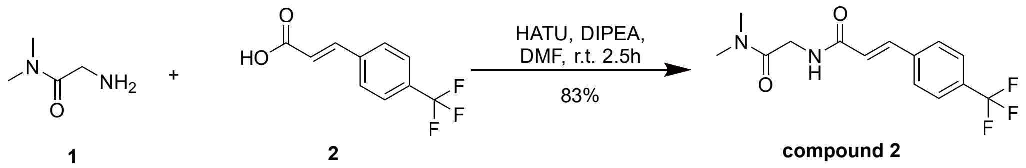


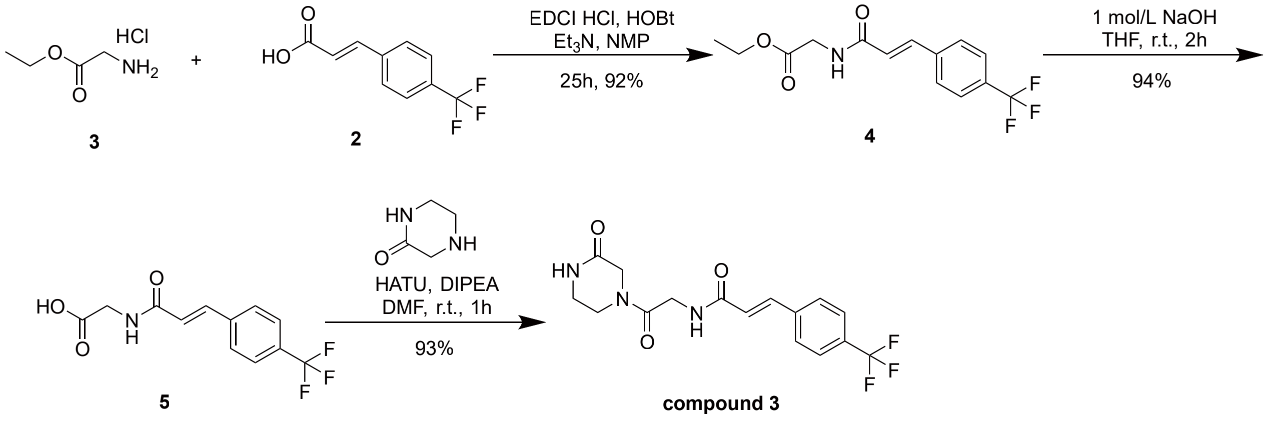


**(E)-N-[2-(dimethylamino)-2-oxo-ethyl]-3-[4-(trifluoromethyl)phenyl]prop-2-enamide (compound 2)**

(E)-3-[4-(trifluoromethyl)phenyl]prop-2-enoic acid (199 mg, 0.92 mmol, 1.00 eq), 2-amino-N,N-dimethyl-acetamide (116 mg, 1.14 mmol, 1.20 eq), DIPEA(0.19 mL, 1.10 mmol, 1.20 eq) and HATU(391 mg, 1.03 mmol, 1.10 eq)were added to DMF (5 mL). The mixture was stirred for 2.5 h at rt. Then, water was added, precipitated solid was separated with filtration. The filtered compound was washed with water and dried over. White solid,

yield 230 mg (0.77 mmol, 83 %).

$^{1}H$ NMR (400 MHz, DMSO-$d_6$): δ [ppm] = 2.87 (s, 3H), 2.98 (s, 3H), 4.08 (d, J = 5.4 Hz, 2H), 7.00 (d, J = 15.8 Hz, 1H), 7.50 (d, J = 15.8 Hz, 1H), 7.75-7.83 (m, 4H), 8.25 (t, J = 5.4 Hz, 1H). $^{13}C$ NMR (101 MHz, DMSO-$d_6$): δ [ppm] = 35.1, 35.7, 40.8, 120.1, 122.8, 124.9, 125.5, 125.8, 128.2, 129.2, 137.2, 139.0, 164.6, 168.0. HRMS (ESI): m/z = 301.1158 (calcd. 301.1157 for $C_{14}H_{16}F_3N_2O_2$ $[M+H]^+$). Purity (HPLC): 99.8 %.

compound 2 $^{1}H$ NMR (400 MHz, $CDCl_3$)

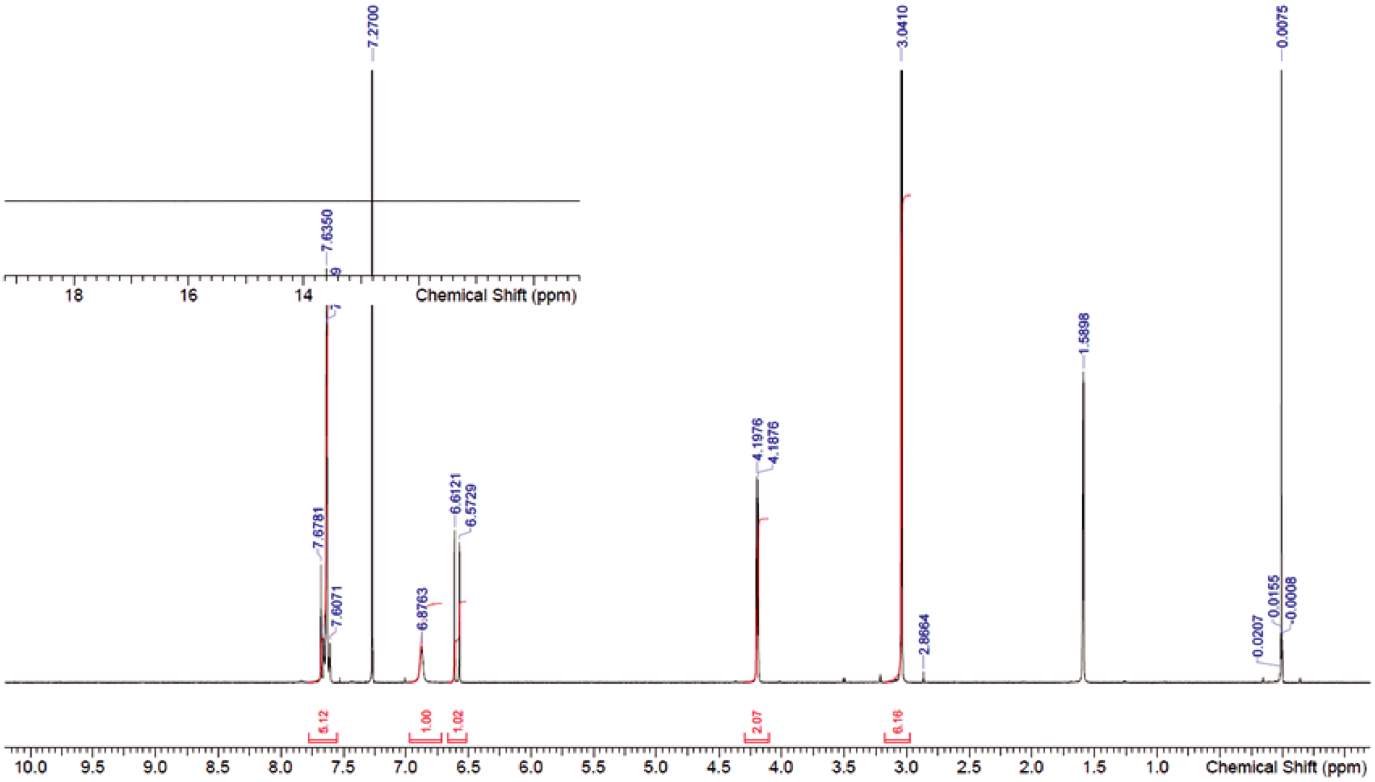


compound 2 $^{13}C$ NMR (101 MHz, DMSO-$d_6$)

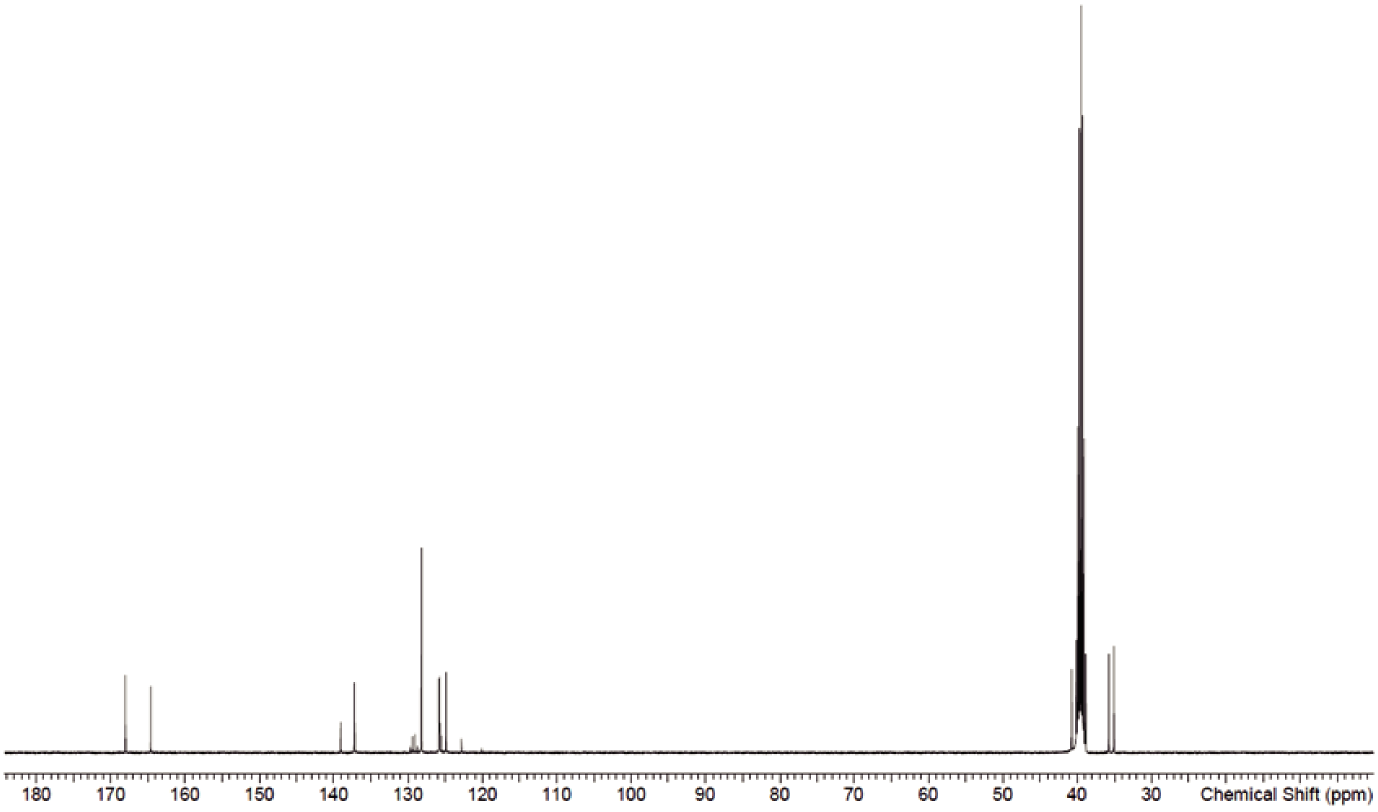


**ethyl 2-[[(E)-3-[4-(trifluoromethyl)phenyl]prop-2-enoyl]amino]acetate (4)**

(E)-3-[4-(trifluoromethyl)phenyl]prop-2-enoic acid (105 g, 485.75 mmol, 1.00 eq), ethyl 2-aminoacetate;hydrochloride (72.1 g, 517 mmol, 1.10 eq), Et3N(150 mL, 1080 mmol, 2.20 eq), WSC HCl (100 g, 521.65 mmol, 1.10 eq) and HOBt (72.2 g, 534 mmol, 2.20 eq) were added to NMP (800 mL). The mixture was stirred for 25 h at rt. Then, water (3 L) was added, precipitated solid was separated with filtration. The filtered compound was washed with water and dried over. White solid,

yield 135 g (448.10 mmol, 92 %).

$^{1}H$ NMR (400 MHz, $CDCl_3$): δ [ppm] = 1.33 (t, J = 7.2 Hz, 3H), 4.20 (d, J = 5.0 Hz, 2H), 4.27 (q, J = 7.2 Hz, 2H), 6.20 (t, J = 5.0 Hz, 1H), 6.55 (d, J = 15.6 Hz, 1H), 7.60-7.68 (m, 4H), 7.68 (d, J = 15.6 Hz, 1H). $^{13}C$ NMR (101 MHz, DMSO-$d_6$): δ [ppm] = 14.4, 41.4, 61.2, 120.4, 123.1, 124.3, 125.8, 126.8, 128.8, 129.9, 138.5, 139.0, 165.8, 170.3. HRMS (ESI): m/z = 302.0999 (calcd. 302.0997 for $C_{14}H_{15}F_3NO_3$ $[M+H]^+$).

ethyl 2-[[(E)-3-[4-(trifluoromethyl)phenyl]prop-2-enoyl]amino]acetate (4) $^{1}H$ NMR (400 MHz, $CDCl_3$)

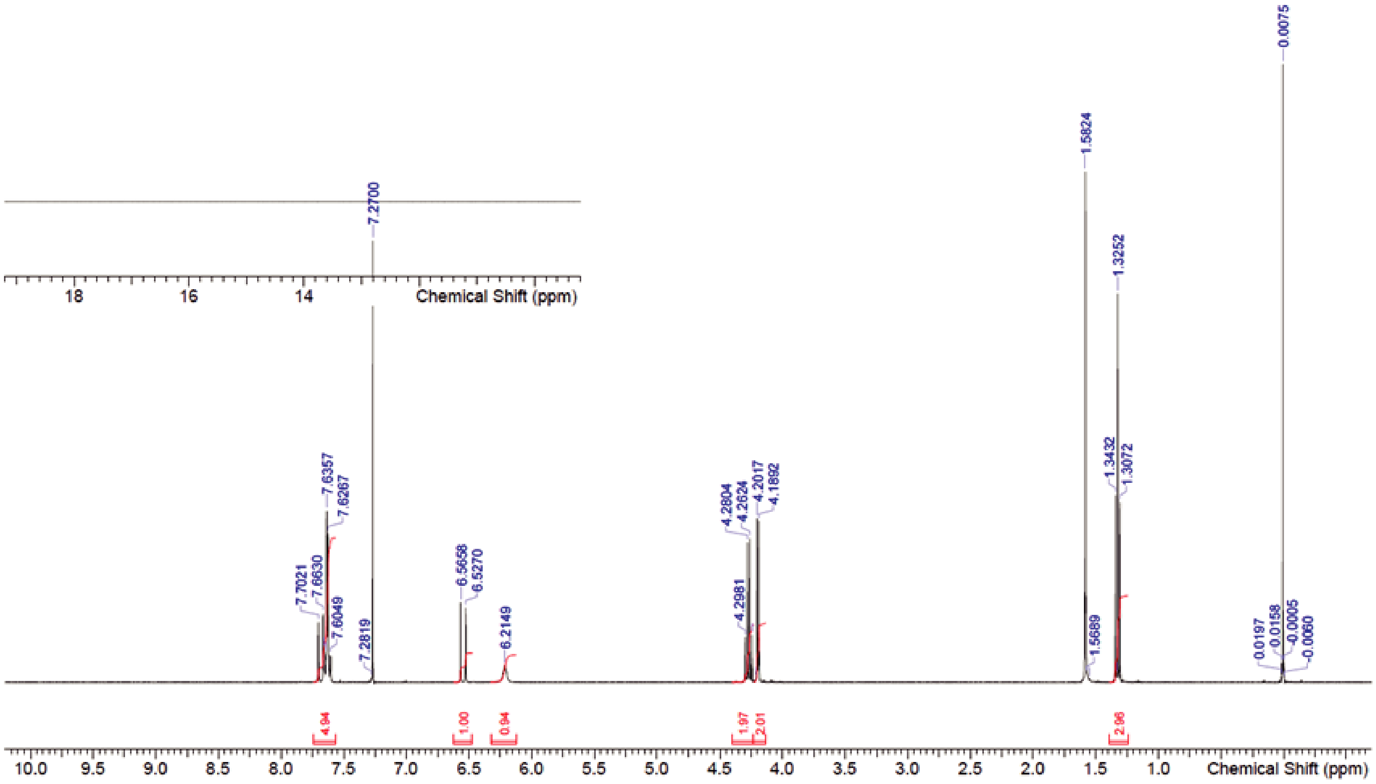


ethyl 2-[[(E)-3-[4-(trifluoromethyl)phenyl]prop-2-enoyl]amino]acetate (4) $^{13}C$ NMR (101 MHz, DMSO-$d_6$)

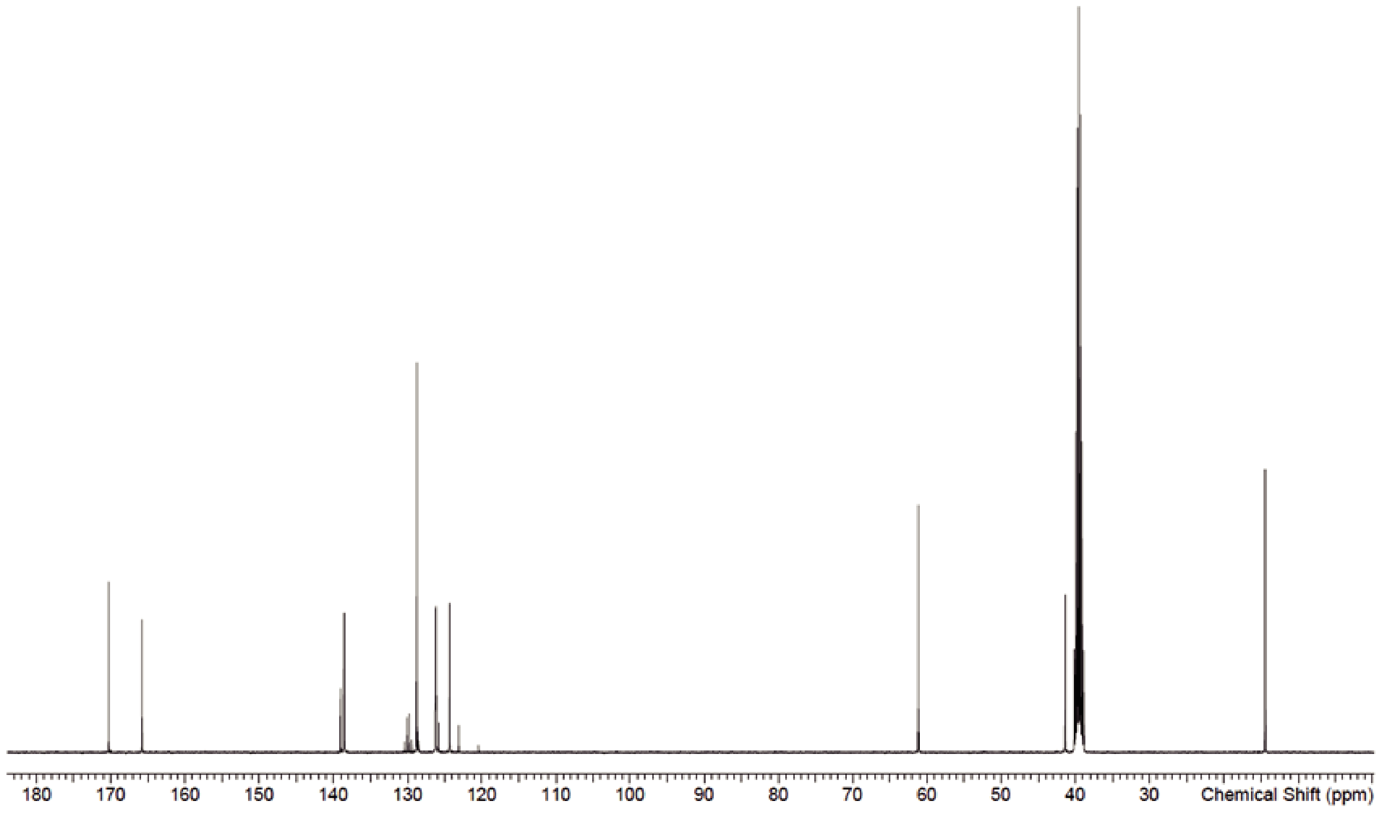


**2-[[(E)-3-[4-(trifluoromethyl)phenyl]prop-2-enoyl]amino]acetic acid (5)**

A mixture of ethyl 2-[[(E)-3-[4-(trifluoromethyl)phenyl]prop-2-enoyl]amino]acetate (40.5 g, 134 mmol, 1.00 eq) and 1 mol/L NaOH(270 mL, 270 mmol, 2.00 eq) in THF (100 mL) was stirred for 2 h at rt. Then 1 mol/L HCl was added to neutralize, precipitated solid was separated with filtration. The filtered compound was washed with water and dried over. White solid,

yield 34.5 g (126 mmol, 94 %).

$^{1}H$ NMR (400 MHz, DMSO-$d_6$): δ [ppm] = 3.91 (d, J = 5.8 Hz, 2H), 6.87 (d, J = 16.0 Hz, 1H), 7.54 (d, J = 16.0 Hz, 1H), 7.74-7.86 (m, 4H), 8.52(t, J = 5.8 Hz, 1H), 12.66 (br s, 1H). $^{13}C$ NMR (101 MHz, DMSO-$d_6$): δ [ppm] = 41.4, 120.4, 123.1, 124.6, 125.1, 126.2, 128.7, 129.9, 138.4, 139.1, 165.7, 171.7. HRMS (ESI): m/z = 274.0686 (calcd. 274.0683 for $C_{12}H_{11}F_3NO_3$ $[M+H]^+$).

2-[[(E)-3-[4-(trifluoromethyl)phenyl]prop-2-enoyl]amino]acetic acid (5) $^{1}$H NMR (400 MHz, DMSO-$d_6$)

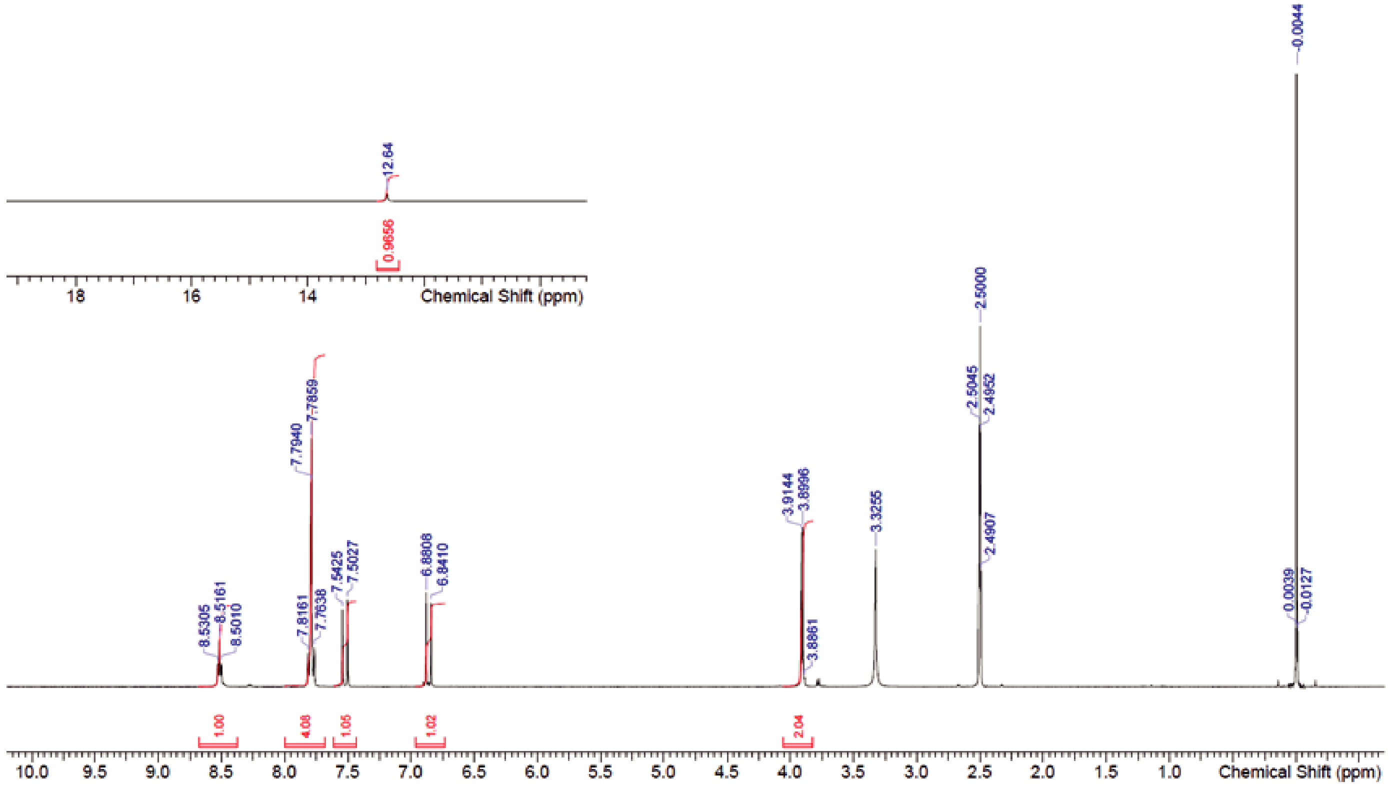


2-[[(E)-3-[4-(trifluoromethyl)phenyl]prop-2-enoyl]amino]acetic acid (5) $^{1}$H NMR (400 MHz, DMSO-$d_6$)

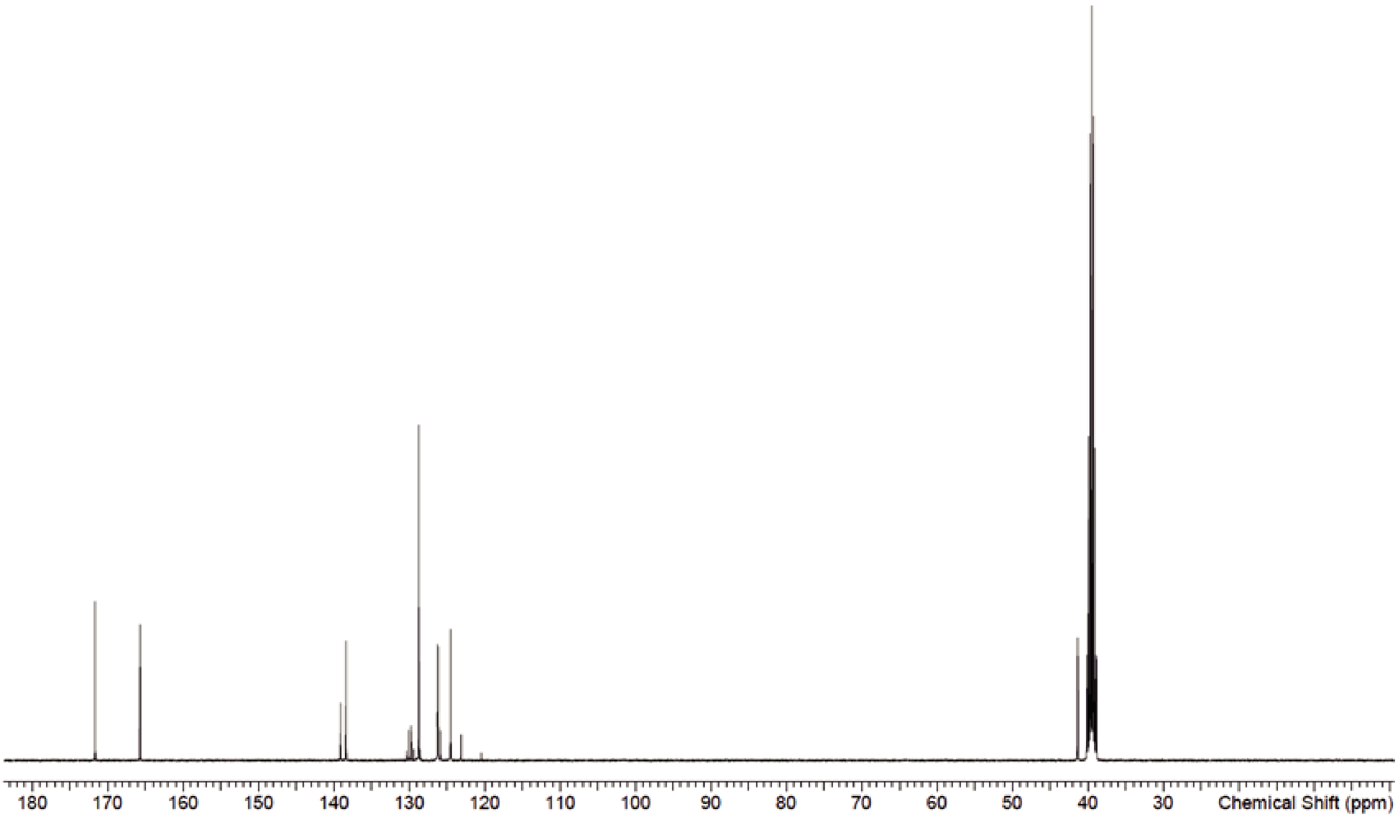


**(E)-N-[2-oxo-2-(3-oxopiperazin-1-yl)ethyl]-3-[4-(trifluoromethyl)phenyl]prop-2-enamide (compound 3)**

2-[[(E)-3-[4-(trifluoromethyl)phenyl]prop-2-enoyl]amino]acetic acid (51.0 mg, 0.19 mmol, 1.00 eq), piperazin-2-one (19 mg , 0.19 mmol, 1.02 eq), DIPEA(0.10 mL, 0.60 mmol, 3.00 eq) and HATU(78 mg, 0.21 mmol, 1.10 eq) were added to DMF (1 mL). The mixture was stirred for 1h at rt. Then, A saturated solution of NH4Cl and CHCl3 were added, the organic layer was separated and the aqueous layer was extracted with CHCl3. The combined organic layers were dried (Na2SO4), filtered and concentrated in vacuo. The crude product was purified by HPLC (column :Xbridge C18, Φ30mm*50mm 5μm, solvent : A: 10mM, Ammonium Carbonate, B: CH3CN, flow rate : 50 ml/min, Gradient20-50% 3min). White solid,
yield 62.0 mg (0.17 mmol, 93 %).

$^{1}$H NMR (400 MHz, DMSO-$d_6$): δ [ppm] = 3.43-3.55 (m, 2H), 3.72 (t, J = 5.6 Hz, 1H, ), 3.90 (t, J = 5.6 Hz, 1H), 4.16 (s, 1H), 4.25 (dd, J = 4.2/12.6 Hz, 2H), 4.32 (s, 1H), 6.21 (br s, 1H), 6.59 (dd, J = 2.0/13.6 Hz, 1H), 6.70-6.85 (br d, 1H), 7.59-7.68 (m, 4H), 7.67 (dd, J = 2.6/13.6 Hz, 1H). $^{13}$C NMR (101 MHz, DMSO-$d_6$): δ [ppm] = 41.3, 41.7, 46.2, 47.7, 120.6, 123.3, 124.7, 126.0, 126.4, 129.0, 130.1, 138.7, 139.2, 166.1, 167.1, 167.9. HRMS (ESI): m/z = 356.1217 (calcd. 356.1215 for $C_{16}H_{17}F_3N_3O_3$ $[M+H]^+$). Purity (HPLC): 96.6 %.

2-[[(E)-3-[4-(trifluoromethyl)phenyl]prop-2-enoyl]amino]acetic acid (5) $^{1}$H NMR (400 MHz, DMSO-$d_6$)

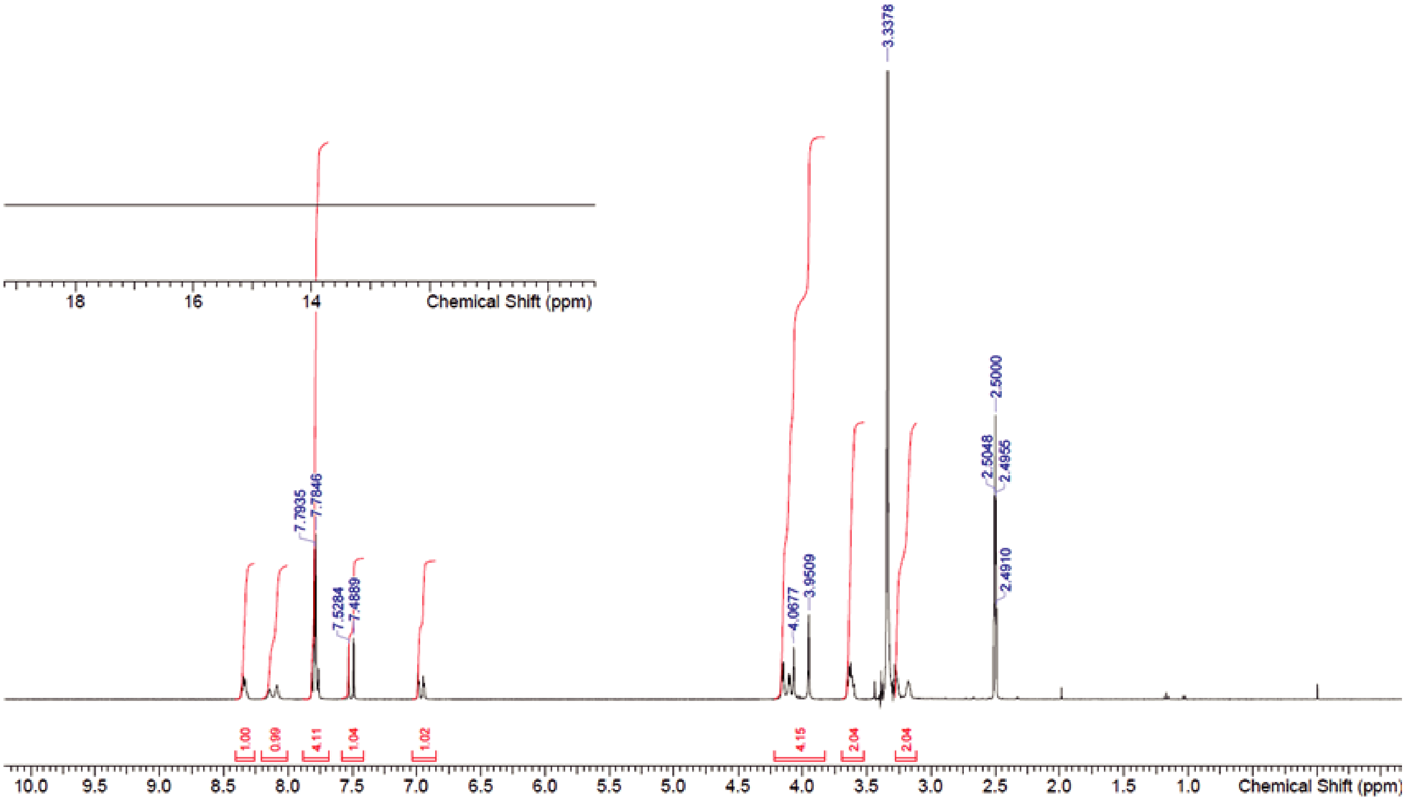


2-[[(E)-3-[4-(trifluoromethyl)phenyl]prop-2-enoyl]amino]acetic acid (5) $^{1}$H NMR (400 MHz, DMSO-$d_6$)

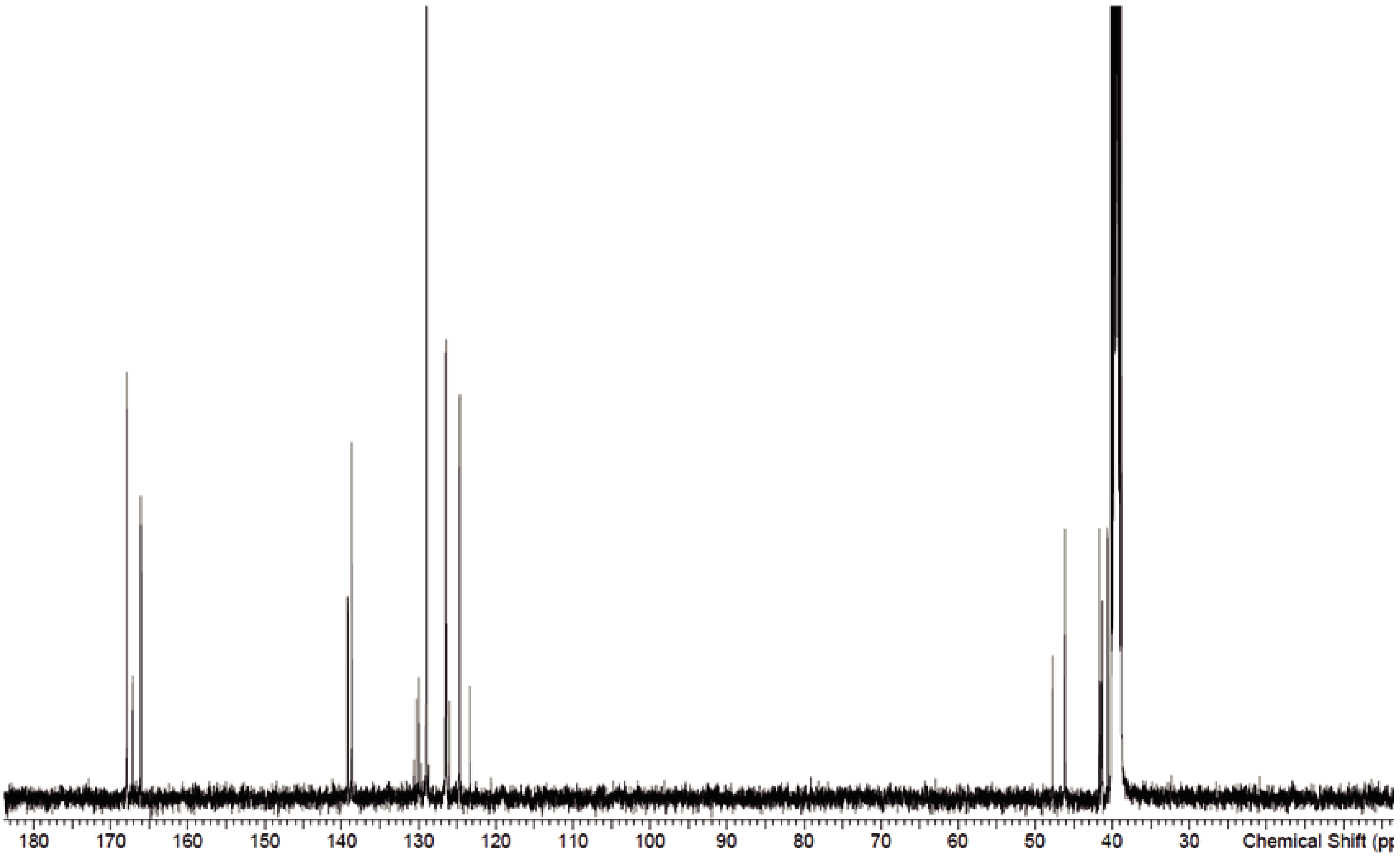
